# Schlafen 11 (SLFN11) overexpression in multiple myeloma and nucleolar translocation in response to bortezomib

**DOI:** 10.64898/2026.02.06.704297

**Authors:** Yasuhiro Arakawa, Daiki Taniyama, Kazuhito Suzuki, Shingo Yano, Yves Pommier

## Abstract

Proteins belonging to the Schlafen family are interferon-inducible and participate in the regulation of antiviral responses, immune signaling and proteotoxic stress. SLFN11 also kills cells with replicative damage, serving as a predictive biomarker for chemotherapeutic response. Here we examined SLFN11 expression and significance in multiple myeloma (MM). The TCGA and MMRF CoMMpass datasets were analyzed for *SLFN11* expression. Bone marrow and cell lines samples were analyzed for SLFN11 protein. SLFN11-knockout MM cell lines were used to explore how SLFN11 affects bortezomib response. Retrospective analysis of the HOVON-65/GMMG-HD4 phase III trial (n=327) assessed clinical relevance. *SLFN11* is consistently highly expressed across MM subtypes (except CD1 and MAF/MAFB) and in normal plasma cells, and its expression strongly correlates with super-enhancer-driven plasma cell transcriptional programs. CD138-positive normal and myeloma plasma cells retain *SLFN11* expression even when proliferative activity (*MKI67*/Ki-67) increases with disease progression. Bortezomib, a first-line MM treatment, induces SLFN11 nucleolar translocation with suppression of ribosomal RNA synthesis. Knocking out SLFN11 in MM cells enhances bortezomib sensitivity and exatecan resistance, supporting SLFN11’s protective role in proteotoxic stress and sensitizing role in replication stress. In the HOVON-65/GMMG-HD4 trial, *SLFN11*-low patients showed selective benefit from bortezomib-based therapy, suggesting that *SLFN11* expression may guide therapeutic stratification in MM.

**Significance:** SLFN11 is highly expressed in normal and malignant plasma cells. Bortezomib induces SLFN11 nucleolar translocation, suppressing ribosomal RNA synthesis and global translation. SLFN11 confers bortezomib resistance while sensitizing to topoisomerase I inhibitors. Clinical analysis supports that low SLFN11 expression predicts bortezomib benefit, providing a mechanistic basis for SLFN11-guided therapeutic stratification.

## Introduction

Multiple myeloma (MM) arises from the clonal malignant proliferation of plasma cells (1). MM molecular subtypes are based on genetic alterations including *MAF*/*MAFB* translocations (MF subtype), *CCND1*/*CCND3* translocations (CD1/CD2 subtypes), MMSET translocations (MS subtype), hyperdiploidy (HY subtype) and proliferation signatures (PR subtype) (2). Comprehensive molecular profiling has expanded molecular understanding, identified high-risk genetic subtypes and elucidated molecular underpinnings of MM (3,4). Despite advances in treatment modalities based on proteasome inhibitors, immunomodulatory drugs and targeted therapies, precision medicine approaches remain limited, highlighting the need to identify additional biomarkers and molecular targets (1,5).

The Schlafen (SLFN) family comprises 10 genes in mice (Slfn1-10) and 6 in humans (6). The human genes (*SLFN* 5, *11*, *12*, *12L*, *13* and *14*) are clustered on chromosome 17q12. In addition to the ribonuclease activity shared by all the Schlafen genes, SLFN11 binds to single-stranded DNA comprising immunostimulant CpG dinucleotides and to the single-stranded protein RPA. SLFN11 acts by irreversibly blocking replication and killing cancer cells in response to DNA damage produced by treatments that create replication stress (7). Thus, SLFN11 is a cell killing mediator in response to replicative DNA damage and an established predictive biomarker for multiple DNA-damaging agents across cancer types (7). Recently, SLFN11 has also been shown to kill cells by inducing TP53-independent apoptosis and blocking ribosome biogenesis (8,9). The predictive value of *SLFN11* expression has been demonstrated in cell line models and across malignancies where high *SLFN11* expression correlates with responses to chemotherapies targeting DNA replication (10–17). Epigenetic inactivation of *SLFN11* occurs in approximately 50% of cancer cell lines and in a large fraction of patient tumors, which leads to chemoresistance (7,12). By contrast, some tumors consistently expressed high *SLFN11* such as acute myeloblastic leukemia (18), mesothelioma (19) and Ewing sarcoma (20) (Figure 1A).

**Figure 1.**
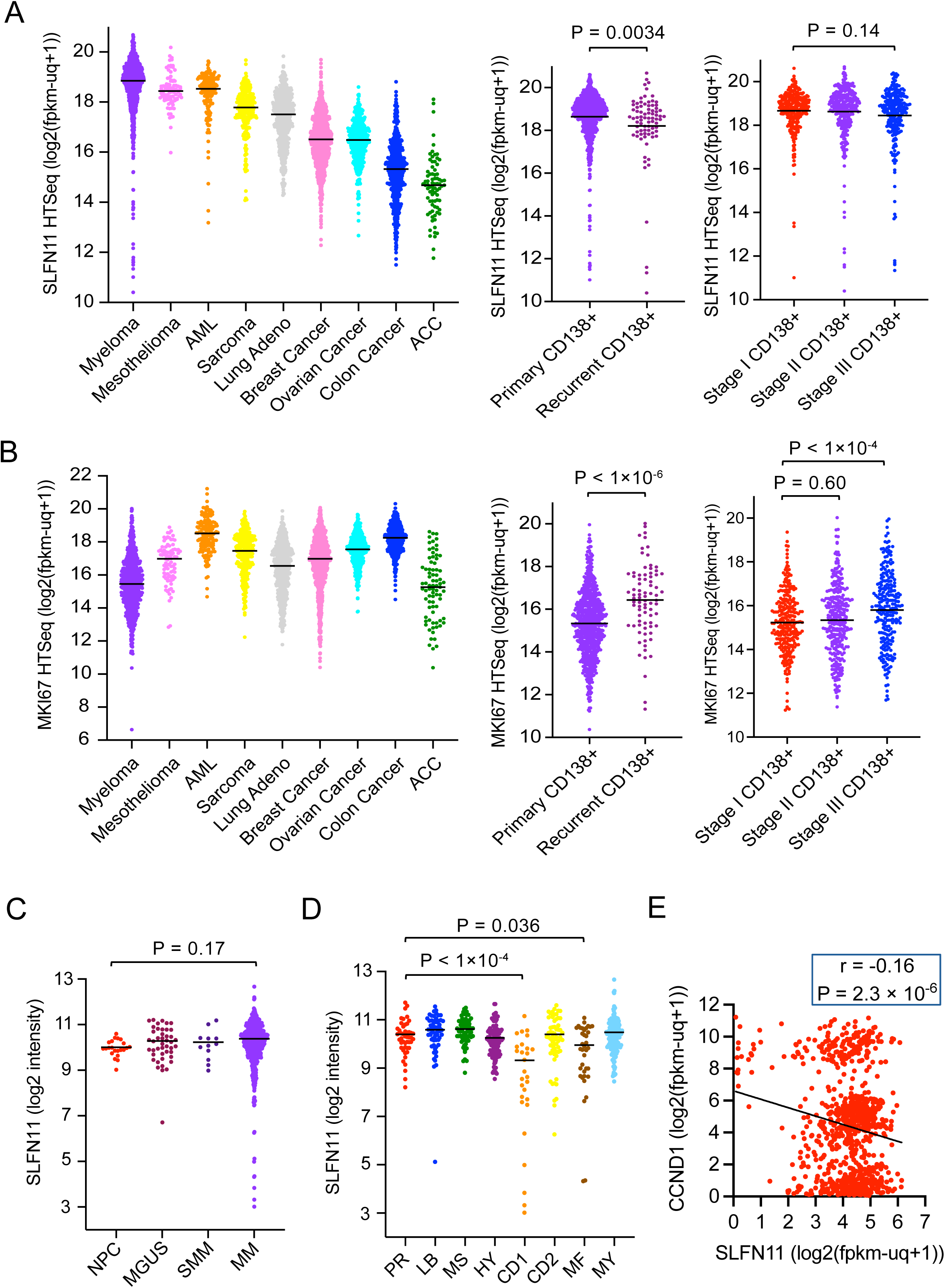
*SLFN11* and *MKI67* expression across cancer types and clinical parameters in multiple myeloma. (A) Left panel: *SLFN11* expression levels (log2(fpkm-uq+1)) across various cancer types from the TCGA and MMRF datasets, with MM showing the highest expression. Middle panel: *SLFN11* expression in primary (newly diagnosed, n = 764) versus recurrent (n = 80) bone marrow-derived CD138+ MM cells (P = 0.0034). Right panel: *SLFN11* expression across disease stages using ISS/R-ISS criteria (P = 0.14). (B) Left panel: *MKI67* expression across cancer types, showing relatively low expression in MM. Middle panel: *MKI67* expression in primary versus recurrent CD138+ cells (P < 1×10⁻□). Right panel: *MKI67* expression across disease stages, with Stage III significantly higher than Stage I (P < 1×10⁻□). (C) *SLFN11* expression (log2 intensity of probe 226743) across plasma cell disorders: normal plasma cells (NPC, n = 22), MGUS (n = 44), smoldering MM (SMM, n = 12), and MM (n = 559) (P = 0.17). This dataset (GSE5900/GSE2658) includes normal controls not available in MMRF. (D) *SLFN11* expression across MM molecular subtypes (n = 559). CD1 and MF subtypes show significantly lower expression compared to PR subtype (P < 1×10⁻□ and P = 0.036, respectively). (E) Correlation between *SLFN11* and *CCND1* expression (r = −0.16, P = 2.3 × 10⁻□). Samples with low *SLFN11* expression (log2(fpkm-uq+1) < 2) predominantly exhibit high *CCND1* expression. In all panels, bars represent mean values.

SLFN11 also act as a regulator of protein translation (8,9), a viral restriction factor (21) and an attenuator of endoplasmic reticulum and proteotoxic stress (22). These functions could have particular significance in malignancies such as MM, which are characterized by high immunoglobulin production and can be prone to inherent proteotoxic stress, which explains the efficacy of proteasome inhibitors as cornerstone therapy of MM.

Because the significance of SLFN11 in MM has until now remained unknown, the present study aimed to characterize *SLFN11* expression across MM subtypes, elucidate SLFN11 molecular interactions and pathway associations, and determine the impact of proteasome inhibition on *SLFN11* expression in MM.

## Methods

### Cell Lines and Patient Specimens

Multiple myeloma cell lines (KMS-27, KMS-34, MM.1S) (23) were obtained from authenticated sources and maintained under standard conditions. Bone marrow specimens from multiple myeloma patients were obtained with institutional ethics approval (Jikei University School of Medicine, approval 37-008). Normal bone marrow tissue arrays (BN29011a and BM241b) were purchased from TissueArray.Com (Derwood, MD, USA).

### CRISPR-Cas9 SLFN11 Knockout and Expression Systems

SLFN11 knockout clones were generated in KMS-27 and KMS-34 cells using CRISPR-Cas9 system with constructs provided by Dr. Junko Murai (12). Doxycycline-inducible *SLFN11* expression was established in U2OS cells via lentiviral transduction.

### Immunohistochemistry and Immunofluorescence

Dual immunohistochemical staining was performed on FFPE bone marrow sections using anti-SLFN11 (Santa Cruz, #sc-515071) and anti-CD138 (Proteintech, #10593-1-AP) antibodies with polymer detection systems. Additional dual staining was performed using anti-CD138 and anti-Ki-67 (Santa Cruz, #sc-101861) antibodies. Immunofluorescence analysis utilized pre-extraction protocols (24) with anti-SLFN11 and anti-Nucleolin antibodies, with imaging performed using Nikon SoRa spinning disk confocal microscopy.

### Ribosomal RNA Synthesis Assay

Nascent RNA synthesis was assessed using 5-ethynyl uridine (EU) incorporation followed by Click-iT RNA Alexa Fluor 488 detection. Signal intensity was quantified with the Fiji software.

### Public Dataset Analysis

*SLFN11* expression was analyzed using MMRF CoMMpass (N=844) and TCGA Pan-Cancer datasets via UCSC Xenabrowser (25,26). Gene Set Enrichment Analysis was performed using blitzGSEA with MSigDB gene sets (27). Samples were stratified by median *SLFN11* expression for pathway analysis. To assess the clinical relevance of SLFN11 expression for bortezomib response, we performed a retrospective subgroup analysis using publicly available gene expression and survival data from the phase III HOVON-65/GMMG-HD4 trial. Details are provided in Supplemental Methods.

### Statistical Analysis

To ensure robust data interpretation, we execute all statistical evaluations through GraphPad Prism 11 software. Comparisons used appropriate tests (t-test, Mann-Whitney U, ANOVA) based on data distribution. P < 0.05 was considered significant. Additional methodological details are provided in the Supplemental Methods.

## Results

### *SLFN11* expression is consistently elevated in multiple myeloma

Analysis of TCGA and MMRF CoMMpass datasets (4,28) revealed that MM express high *SLFN11* transcript levels, exceeding mesothelioma, sarcoma, and acute myeloid leukemia (AML) (Figure 1A, left) (16,18–20). Only 2.8% show low *SLFN11* expression (log2(fpkm-uq+1) < 16) (Figure 1A).

Examination of different disease stages and conditions shows that *SLFN11* expression is statistically lower in recurrent bone marrow samples than in samples from primary disease (P = 0.0034) (Figure 1A, middle). However, the magnitude of this difference is relatively small, suggesting that *SLFN11* expression remains high despite disease recurrence. No significant variation in *SLFN11* expression was observed across different disease stages (P = 0.14) (Figure 1A, right).

We examined *MKI67* expression, as both SLFN11 and proliferation status predict sensitivity to DNA-damaging agents (29,30). Multiple myeloma typically exhibits relatively low proliferative activity, which may contribute to conventional chemotherapy resistance (31). *MKI67* expression was notably lower in MM than other cancer types (Figure 1B, left). Yet, *MKI67* was significantly elevated in recurrent versus primary samples (P < 1×10^-6^, >2-fold), and Stage III versus Stage I (P < 1×10^-4^) (Figure 1B, middle and right panels).

To explore *SLFN11* expression during plasma cell disorder progression, we analyzed *SLFN11* levels in normal plasma cells (NPC), monoclonal gammopathy of undetermined significance (MGUS), smoldering multiple myeloma (SMM) and symptomatic MM using published gene expression microarray data (GSE5900 and GSE2658) (32). No statistically significant difference was observed between these groups (P = 0.17) (Figure 1C), indicating that normal plasma cells inherently express high levels of *SLFN11*, and that this elevated expression is maintained throughout malignant transformation and progression to symptomatic MM.

Analysis of 1,019 cancer cell lines in public databases (https://discover.nci.nih.gov/cellminercdb/) revealed that approximately one-third of MM cell lines exhibit *SLFN11* deficiency and nearly half show high *MKI67* expression (Figure S1A), contrasting with the consistently high *SLFN11* expression (97.2%) and low proliferative activity observed in patient-derived MM samples. This difference is likely due to the fact that cell line selection favors clones with high proliferation potential.

### *SLFN11* expression across MM molecular subtypes

We next explored whether *SLFN11* expression differs across the established MM molecular subtypes. Analysis of the microarray dataset GSE2658 (2) revealed variation in *SLFN11* expression among the molecular subgroups (Figure 1D). Both the CD1 and MAF/MAFB (MF) subtypes exhibit significantly lower *SLFN11* expression compared to the proliferation (PR) subtype (P < 1×10^-4^ and P = 0.036, respectively). Accordingly, samples with low *SLFN11* (log2 intensity < 7) were predominantly classified as CD1 subtype. No significant difference in *SLFN11* expression was observed between the PR subtype and other molecular subtypes: MMSET (MS), hyperdiploid (HY), low bone disease (LB) and CD2 (Figure 1D).

As dysregulation of D-type cyclins is a known driver of MM pathogenesis (2), we examined cyclin D expression in the MMRF CoMMpass dataset. Samples with low *SLFN11* expression (log2(fpkm-uq+1) < 2) demonstrated high expression of *CCND1* (Figure 1E). While not all *CCND1*-high samples showed low *SLFN11* expression, the inverse relationship suggests that *CCND1* overexpression is associated with *SLFN11* suppression in a distinct subset of MM.

Additional analysis of the MMRF CoMMpass dataset across the recently defined RNA expression subtypes revealed that a subset of samples in the CD subtypes (CD1, CD2a, CD2b) exhibit low *SLFN11* expression. We also observed notable differences in *SLFN11* expression among hyperdiploid (HRD) samples, varying by their specific additional genetic alterations (Figure S1B). Yet, copy number-based subtypes showed minimal variation in *SLFN11* expression (Figure S1C), suggesting that high *SLFN11* expression in MM is primarily driven by transcriptional upregulation.

### CD138-positive myeloma cells express SLFN11 in patient bone marrow samples

To determine SLFN11 protein expression in MM patient bone marrow samples, we performed double immunocytochemical staining of SLFN11 and CD138, an established marker routinely used to identify malignant cells in MM. Figure 2A shows that CD138-positive myeloma cells (81-98%) co-express SLFN11. Staining patterns, both of primary (P) and relapsed (R) specimens revealed that SLFN11 (brown) is mainly localized to the nucleus while CD138 (blue) shows characteristic membrane staining.

**Figure 2.**
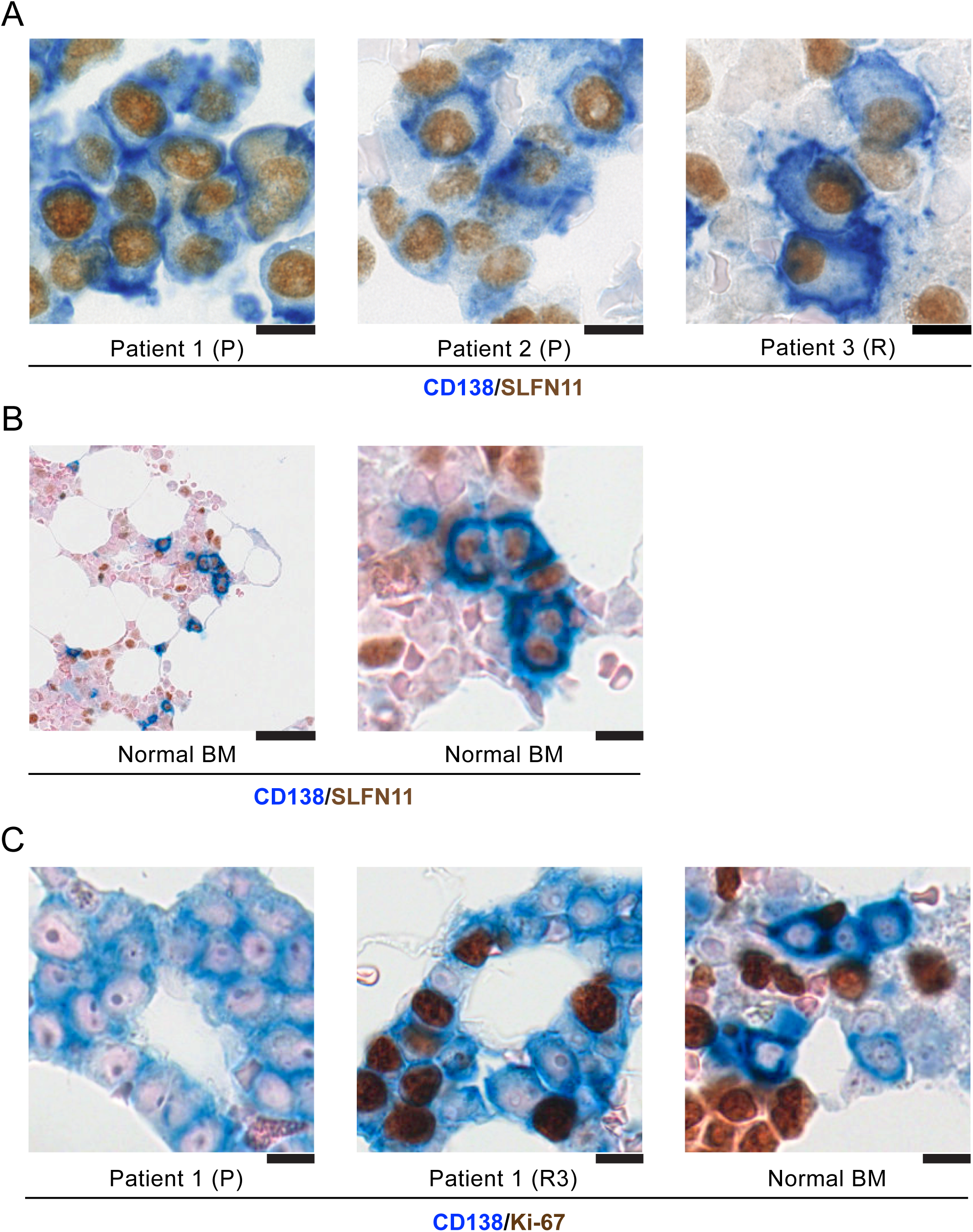
Expression of SLFN11, CD138 and Ki-67 in multiple myeloma patient samples and normal bone marrow. (A) Representative dual immunohistochemical staining for CD138 (blue, membrane) and SLFN11 (brown, nuclear) in bone marrow samples from multiple myeloma patients. Patient 1 and Patient 2 are primary (P) (newly diagnosed), while Patient 3 (R) represents a sample from a recurrent patient. 81-98% CD138-positive myeloma cells express SLFN11. (B) Dual immunohistochemical staining for CD138 (blue) and SLFN11 (brown) in normal bone marrow. Left panel shows low magnification and right panel high magnification of the same field. Normal bone marrow contains 5.5-6.8% plasma cells, of which 72-85% demonstrate SLFN11 positivity, indicating that *SLFN11* expression is an intrinsic feature of normal plasma cells. Scale bars: 50 μm (left) and 10 μm (right). (C) Representative dual immunohistochemical staining for CD138 (blue) and Ki-67 (brown, nuclear) to assess proliferative activity. Left panel and middle panels show Patient 1 at primary diagnosis (P) and third relapse (R3), respectively. Right panel shows normal bone marrow. The plasma cell proliferation index (PCPI), defined as the percentage of CD138-positive cells co-expressing Ki-67, is markedly elevated in myeloma specimens compared to normal bone marrow, with further increases observed at relapse. Scale bars: 10 μm.

Examination of normal bone marrow samples containing plasma cells representing 5-7% of total bone marrow cells, demonstrated that normal plasma cells are generally SLFN11 positive (72-85%; Figure 2B). These findings demonstrate that SLFN11 expression is an intrinsic feature of both normal and malignant plasma cells.

To assess single-cell proliferative activity, we performed CD138/Ki-67 double staining and calculated the plasma cell proliferation index (PCPI), defined as the percentage of CD138-positive cells co-expressing Ki-67 (33). Consistent with our transcriptome findings (Figure 1B), comparison of MM patient samples with normal bone marrow demonstrated elevated PCPI in myeloma specimens, with further increase observed in relapsed samples (Figure 2C).

Quantitative analysis of four distinct cell populations based on SLFN11/CD138 expression patterns (CD138+/SLFN11+, CD138+/SLFN11-, CD138-/SLFN11+, and CD138-/SLFN11-) was performed on sequential bone marrow samples from three MM patients and normal bone marrow controls (Figure S2A, B). Although the total percentage of CD138-positive myeloma cells fluctuated over time with disease progression and treatment response, most myeloma cells consistently maintained SLFN11 expression throughout the disease course. The PCPI analysis confirmed progressive increases with each relapse across all three patients, though the absolute values remained relatively modest in patients 2 and 3 compared to patient 1. In contrast, normal bone marrow samples showed minimal PCPI values (Figure S2C).

Taken together, these observations establish SLFN11 expression as a feature of CD138-positive myeloma cells, which is maintained from normal plasma cells through malignant transformation and MM disease progression.

### SLFN11 is expressed with plasma cell differentiation markers

To explore the relationship between SLFN11 and known plasma cell markers, we analyzed the MMRF CoMMpass dataset, where clinical samples are enriched for plasma cells through CD138 antibody-based magnetic bead selection prior to RNA sequencing. Significant positive correlations were observed between *SLFN11* expression and the expression of established plasma cell differentiation markers: *CD138* (*SDC1* HUGO name), *CD38* and *BCMA* (TNF Receptor Superfamily Member 17; *TNFRSF17*) (Figure 3A).

**Figure 3.**
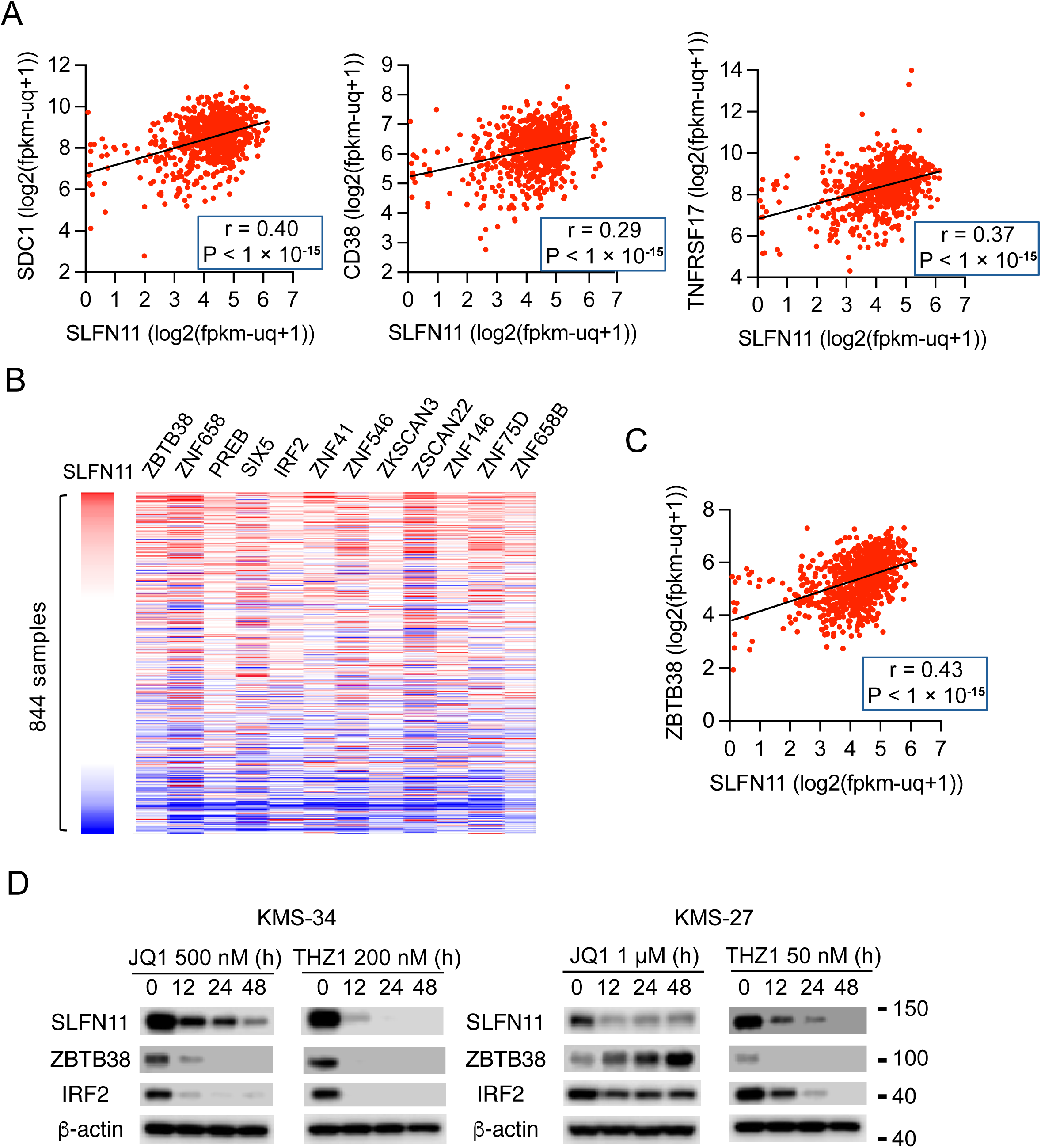
Correlations between the expression of *SLFN11* and plasma cell markers, and *SLFN11* transcriptional regulation in multiple myeloma. (A) Scatter plots demonstrating positive correlations between *SLFN11* expression and plasma cell markers (n = 844 multiple myeloma samples from the MMRF CoMMpass Study. (B) Heatmap displaying the mRNA expression of the top 12 transcription factors and epigenetic enzymes with the strongest correlation with *SLFN11* expression (samples are from the MMRF CoMMpass Study. Red indicates high expression; blue low expression. (C) Scatter plot highlighting the positive correlation between *SLFN11* and *ZBTB38* expression. (D) Representative Western blots showing time-dependent effects of super-enhancer inhibitors on SLFN11, ZBTB38, and IRF2 protein levels in MM cell lines. Left panels show KMS-34 cells treated with JQ1 (BET inhibitor, 500 nM) or THZ1 (CDK7 inhibitor, 200 nM) for 0, 12, 24, and 48 hours. Right panels show KMS-27 cells treated with JQ1 (1 μM) or THZ1 (50 nM) for the same time points. β-actin serves as a loading control.

In contrast, *SLFN11* expression showed no significant correlation with markers of other hematopoietic lineages, including *CD8A* (T-cell marker), *CD79A* (B-cell marker), or *CD68* (monocyte/macrophage marker) (Figure S3A). This specificity confirms that the high *SLFN11* expression observed in MM is associated with the plasma cell differentiation program rather than contaminating immune cell populations.

### Transcriptional regulation of SLFN11 in multiple myeloma

To identify potential transcriptional regulators of *SLFN11* in multiple myeloma, we examined the correlations between *SLFN11* expression and the expression of 122 transcription factors and epigenetic enzymes involved in plasma cell differentiation (Figure S3B) (34). The 12 factors showing strongest correlation with *SLFN11* are presented in Figure 3B, with *ZBTB38* most highly correlated (r = 0.43, Figure 3C). This result is consistent with the coordinated regulation of *SLFN11* with the plasma cell transcriptional network.

Since super-enhancers have been shown to regulate key transcription factors essential for multiple myeloma cell state, including *IRF4*, *PRDM1*, *MYC*, and *XBP1* (35), we treated two MM cell lines (KMS-34 and KMS-27) with inhibitors of super-enhancer-mediated gene expression. Both JQ1 (BET inhibitor) and THZ1 (CDK7 inhibitor), which target super-enhancer-mediated transcriptional control (36) reduced SLFN11 protein levels in a time-dependent manner, along with concomitant decrease in IRF2 (Figure 3D), indicating that SLFN11 expression is under super-enhancer control in MM.

By contrast, *FLI1* expression, a known transcriptional regulator of *SLFN11* in Ewing sarcoma (20) and leukemia (18), showed only a weak and non-significant correlation with *SLFN11* expression in MM (r = 0.15; Figure S3C), implying context-specific regulatory mechanisms for *SLFN11* expression (20). These results establish a strong association between *SLFN11* expression and the plasma cell differentiation super-enhancer program in MM.

### Molecular pathway signatures associated with *SLFN11* expression in MM

To gain insights into the functional implications of *SLFN11* expression in MM, we performed pathway analyses comparing the samples with the highest and lowest *SLFN11* expression. Gene Ontology (GO) enrichment analysis and heatmap visualization of differentially expressed genes across 844 MM samples reveal distinct biological processes associated with *SLFN11* expression in MM (Figure 4A). *SLFN11*-high samples showed significant enrichment in pathways related to endoplasmic reticulum (ER) stress, unfolded protein response (UPR), ER-associated degradation (ERAD) and Golgi transport mechanisms. These pathways are critical for plasma cells to manage their high immunoglobulin secretory burden. In contrast, GO analysis revealed that *SLFN11*-low samples preferentially express genes related to ubiquitin-protein ligase activity, ubiquitin transferase functions and ribosomal protein transcription, suggesting interactions between SLFN11 and protein quality control mechanisms (8,9).

**Figure 4.**
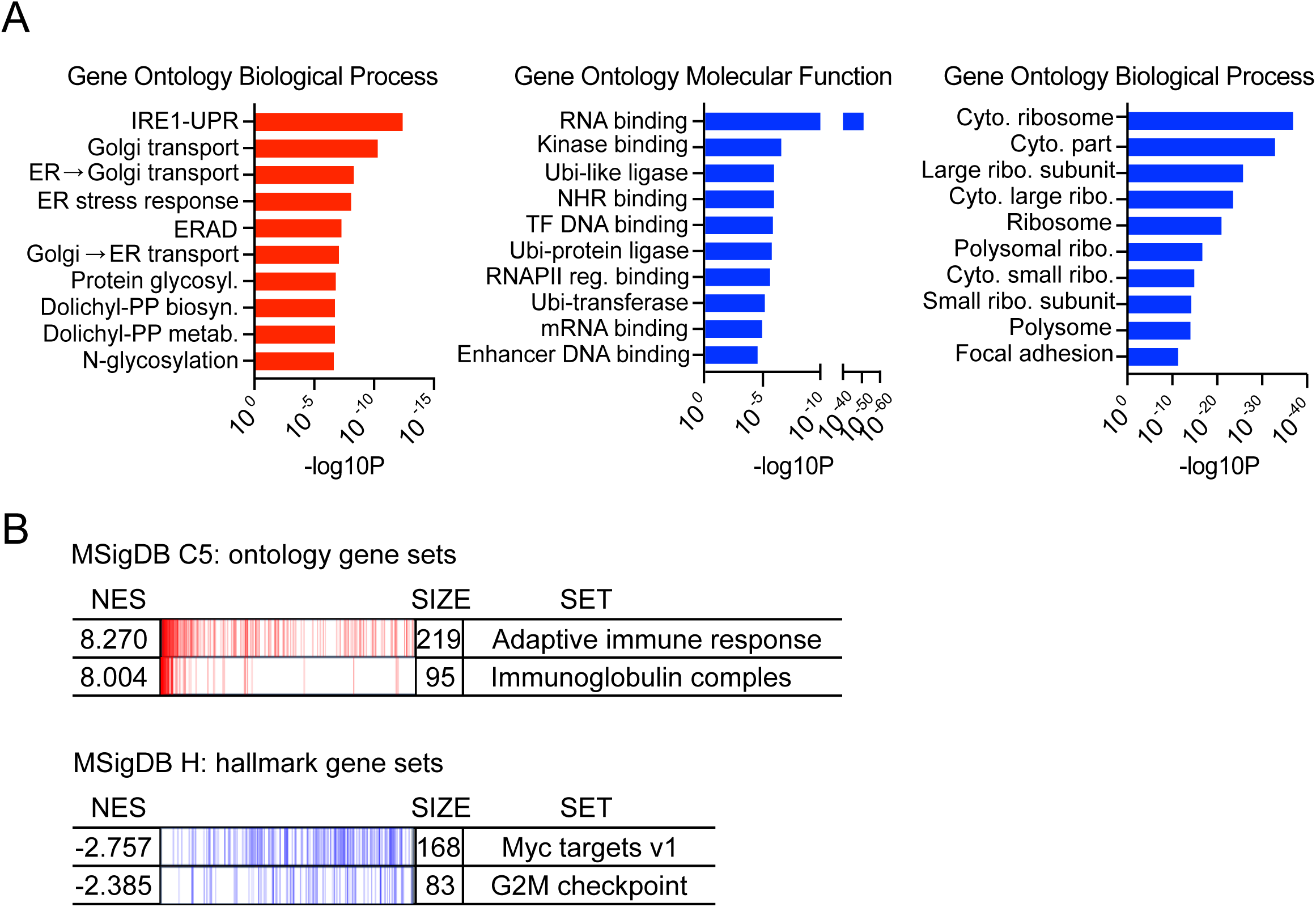
Pathway analysis of *SLFN11*-associated genes in multiple myeloma. (A) Gene Ontology (GO) enrichment analysis comparing MM samples with the highest and lowest *SLFN11* expression (n = 844, divided by median expression). Red bars indicate pathways upregulated in *SLFN11*-high samples, while blue bars show pathways enriched in *SLFN11*-low samples. Left panel: biological processes; middle panel: molecular functions; right panel: cellular components. The x-axis represents -log10(P-value). (B) Gene Set Enrichment Analysis (GSEA) comparing *SLFN11*-high versus *SLFN11*-low samples. Upper panel: MSigDB C5 ontology gene sets showing positive enrichment in *SLFN11*-high samples, including adaptive immune response (NES = 8.270) and immunoglobulin complex (NES = 8.004). Lower panel: MSigDB Hallmark gene sets showing negative enrichment in *SLFN11*-high samples, including MYC targets v1 (NES = −2.757) and G2M checkpoint (NES = −2.385). NES: Normalized Enrichment Score.

Gene Set Enrichment Analysis (GSEA) provides additional insights into the molecular signatures associated with *SLFN11* expression (Figure 4B). *SLFN11*-high samples show significant positive enrichment in adaptive immune response, immunoglobulin complex, and antigen binding gene sets. Conversely, samples with the lowest *SLFN11* expression exhibit enrichment in key proliferation-related pathways such as G2M checkpoint and MYC targets.

Heatmap visualization of the differentially expressed genes across 844 MM samples illustrates the coordination between *SLFN11* expression and genes involved in ER stress response, ERAD, ubiquitin-protein ligase activity, ubiquitin transferase functions, and ribosomal components (Figure S4A). More comprehensive GSEA analysis reveals that *SLFN11*-low samples are enriched in pathways related to TNFα signaling via NFκB, inflammatory response, apoptosis and proliferation markers, while samples with the highest *SLFN11* expression show positive enrichment in cytoplasmic translation, ribosome function, and lymphocyte-mediated immunity (Figure S4B).

Collectively, these findings suggest that *SLFN11* expression in MM is associated with plasma cell phenotype characterized by high secretory activity and ER stress response, while *SLFN11*-low samples exhibit features of increased proliferation and inflammatory signaling.

### SLFN11 translocation to nucleoli upon bortezomib treatment

Because proteasome inhibitors are used as first line therapy in multiple myeloma, we examined the cellular distribution of SLFN11 in MM cell lines treated with the proteasome inhibitor, bortezomib. To select the MM cell lines with high SLFN11 expression, we used the CellMinerCDB website (https://discover.nci.nih.gov/cellminercdb/) (37) to analyze the Broad Institute cell line genomic and drug response database (Figure 5A). Immunoblot analysis confirmed SLFN11 protein expression in the MM cell lines MM.1S, KMS-34, KMS-27, while the osteosarcoma cell line U2OS, known to lack SLFN11 expression (38), served as a negative control (Figure 5B).

**Figure 5.**
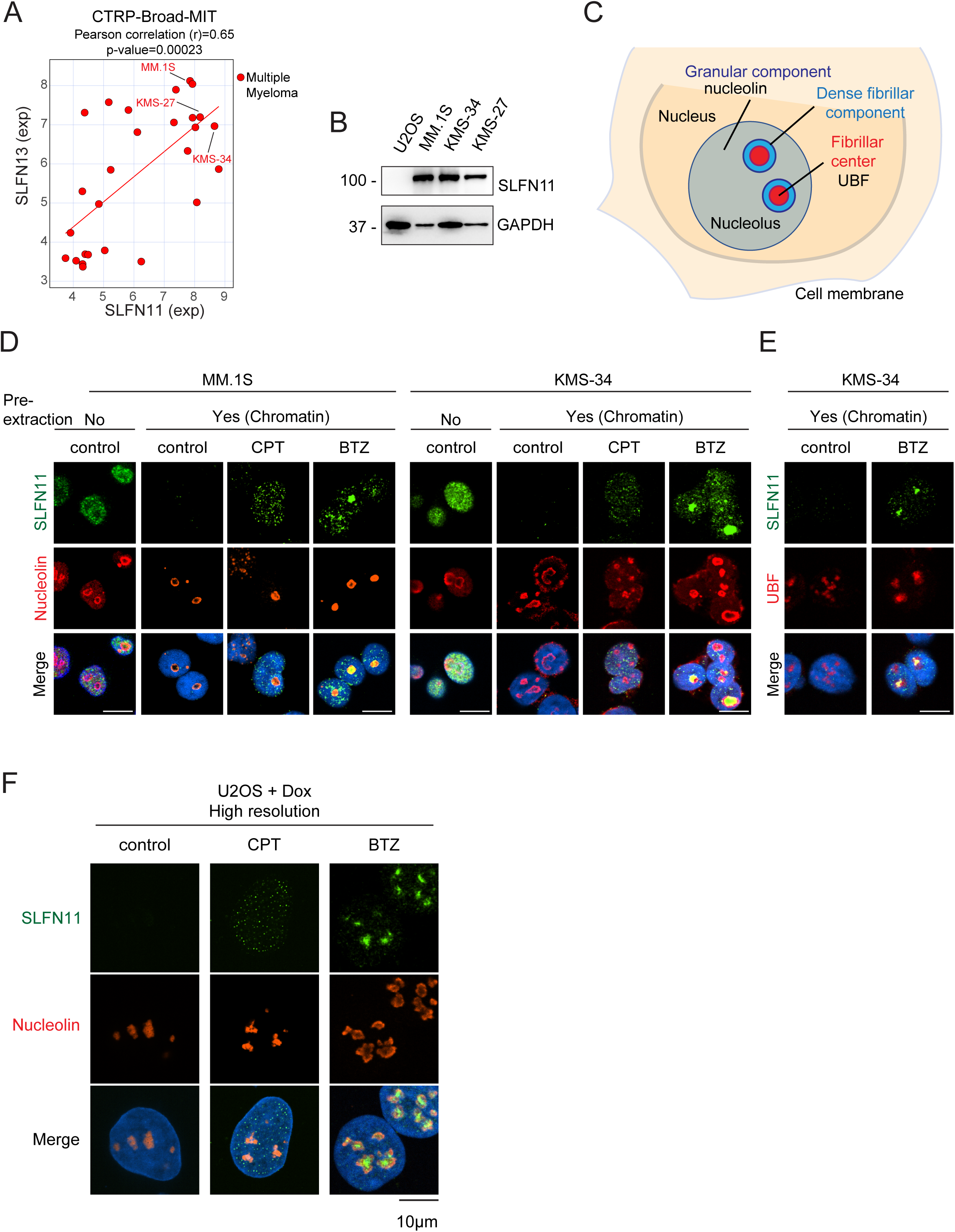
SLFN11 expression in multiple myeloma cell lines, and SLFN11 translocation to nucleoli by bortezomib. (A) Correlation between the expression of *SLFN11* and *SLFN13* in CTRP-Brad-MIT cell line database (https://discover.nci.nih.gov/cellminercdb/). Correlation coefficient and P value are indicated above the figure. (B) Representative SLFN11 immunoblots in MM.1S, KMS-34, and KMS-27 cell lines. Human osteosarcoma cell line U2OS was used as negative control. (C) Scheme of the nucleolus comprising the fibrillar center, the dense fibrillar component, and the granular component. (D) Representative immunofluorescence images showing total SLFN11 (green, left, no pre-extraction) and chromatin-bound SLFN11 (green, right) with nucleolin (red), and DAPI (blue) in MM.1S and KMS-34 cells after 4-hour treatment with bortezomib [BTZ, 0.5µM (MM.1S), 5 µM (KMS-34)] or camptothecin [CPT, 1µM (MM.1S), 5 µM (KMS-34)]. Scale bar: 10 µm. (E) Representative immunofluorescence images showing total chromatin-bound SLFN11 (green) with UBF (red), and DAPI (blue) in KMS-34 cells after 4-hour treatment. Scale bar: 10 µm. (F) Representative immunofluorescence image showing BTZ-induced SLFN11 nucleolar translocation in U2OS with doxycycline-inducible SLFN11 expression. Cells were treated with CPT or BTZ (1 µM) for 4 hours. Right: Quantification of SLFN11 signal intensities in individual cells (n = 2453-2927 cells per condition). ***P < 0.001, ****P < 0.0001 (one-way ANOVA). a.u., arbitrary units.

Given the recent findings linking SLFN11 and nucleolar functions (9), we used established markers to identify the distinct nucleolar regions: UBF (Upstream Binding Factor) for the fibrillar center and nucleolin for the granular component (Figure 5C). Standard immunofluorescence of untreated MM.1S and KMS-34 cells showed the expected predominantly diffuse SLFN11 staining (green) in the nucleus. The SLFN11 staining of untreated cells was largely excluded from nucleoli in MM.1S cells, as evidenced by the minimal overlap with nucleolin (red), though variable nucleolar overlap was observed in KMS-34 cells (Figure 5D, left). This pattern recapitulates the SLFN11 distribution observed in untreated bone marrow samples (see Figure 2A).

As expected under base-line conditions, upon mild detergent pre-extraction, SLFN11 was no longer detectable due to its known loose chromatin binding (39). By contrast following treatment with camptothecin (CPT), a topoisomerase I poison known to induce replicative DNA damage, SLFN11 chromatin foci were readily detectable throughout the nuclei without enrichment in nucleoli (Figures 5D and S5C) (17,39). Unexpectedly, treatment with bortezomib only induced intense SLFN11 staining in nucleoli both in the MM.1S and KMS-34 cells, as evidenced by increased co-localization of SLFN11 with nucleolin and UBF (Figures 5E and S5D). Bortezomib-induced SLFN11 nucleolar translocation was also observed with DOX-inducible SLFN11-expressing osteosarcoma U2OS cells (Figure 5F) implying that this effect is not limited to MM cells.

### SLFN11 suppresses ribosomal RNA synthesis in response to bortezomib

Because the bortezomib-induced tight binding of SLFN11 to the fibrillar center of nucleoli suggested a potential role for SLFN11 in regulating nucleolar functions in response to proteotoxic stress, we investigated the functional significance of the nucleolar translocation of SLFN11 in response to bortezomib. To do so, we generated SLFN11 knockout (KO) clones from the KMS-34 and KMS-27 cell lines using the CRISPR-Cas9 technology (12,39). Knockout was confirmed by immunofluorescence and immunoblot analyses (Figure 6A-B). To assess the impact of SLFN11 on ribosomal RNA (rRNA) synthesis (9), we performed 5-ethynyl uridine (EU) incorporation assays (9). In untreated conditions, both wild-type and SLFN11 KO cells showed robust EU signals, reflecting active RNA synthesis. Following bortezomib treatment, EU incorporation was significantly reduced in wild-type cells, whereas SLFN11 KO cells maintained high levels of EU incorporation (Figure 6C). Quantitative analysis confirmed the significant difference in EU signals between wild-type and SLFN11 KO cells after bortezomib treatment (p < 0.0001) (Figure 6D).

**Figure 6.**
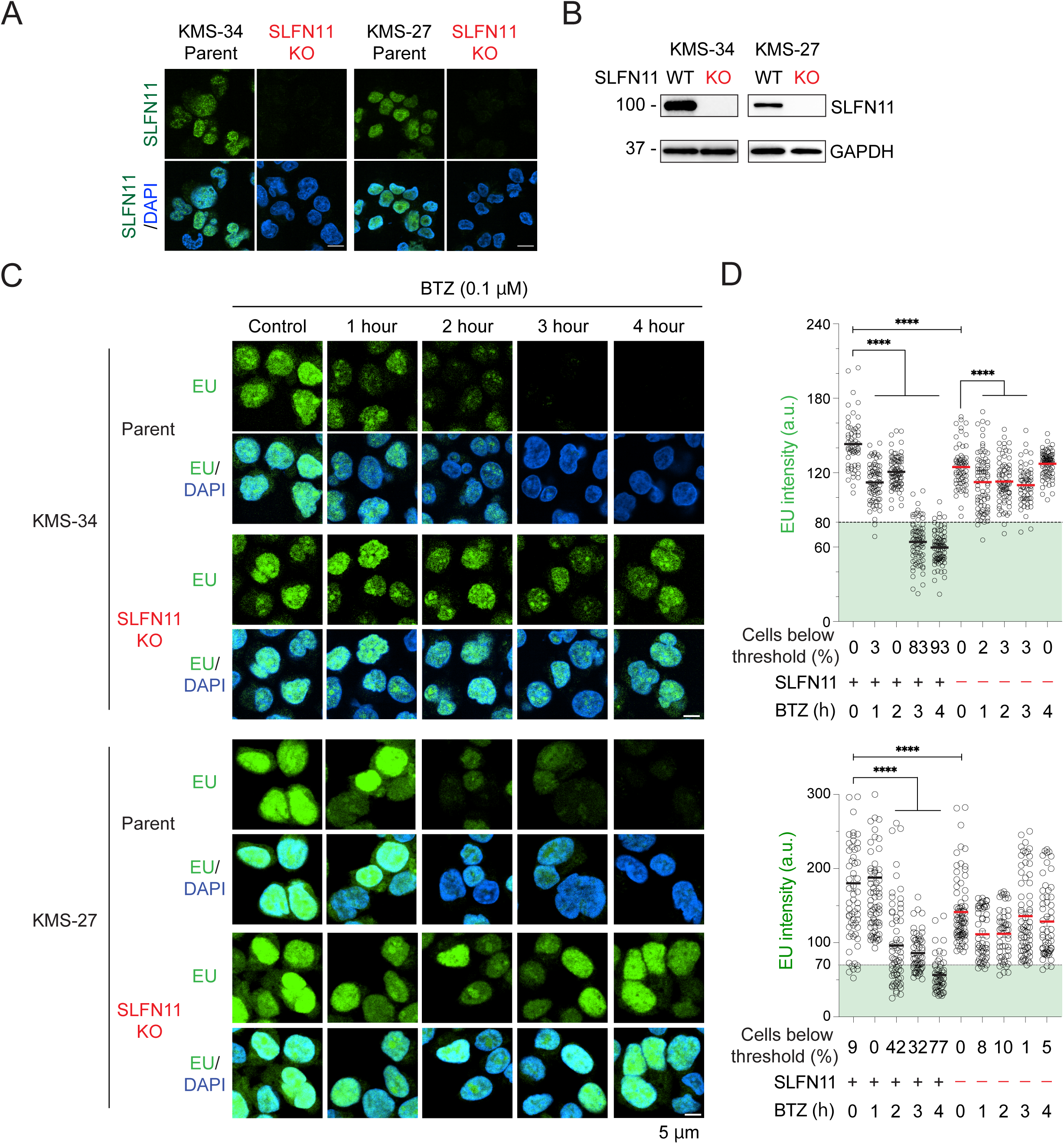
SLFN11-dependent suppression of ribosomal RNA synthesis by bortezomib. (A) Representative immunofluorescence images showing total SLFN11 (green, no pre-extraction) and DAPI (blue) in parental and SLFN11 knockout (KO) KMS-34 cell line. Scale bar: 10 µm. (B) SLFN11 immunoblots for whole-cell lysates from wild-type (WT) and SLFN11 KO clones of KMS-34 cells. (C) Representative immunofluorescence images showing 5-ethynyluridine (EU) incorporation (green) and DAPI (blue). Upper panels: KMS-34 cells. Lower panels: KMS-27 cells. WT and SLFN11 KO clones were treated with BTZ (0.1 µM) for indicated timepoints. Scale bar: 5 µm. (D) Quantification of EU signal intensities in individual cells. Mean ± SEM are shown. Upper panel: KMS-34 cells (threshold = 80). Lower panel: KMS-27 cells (threshold = 70). Percentages of cells below threshold are indicated. ****P < 0.0001 (one-way ANOVA). a.u., arbitrary units.

To further establish the role of SLFN11 in suppressing rRNA synthesis under proteotoxic stress, we employed DOX-inducible SLFN11-expressing U2OS cells (Figure S6A). EU incorporation assays demonstrated that induction of *SLFN11* expression significantly suppresses rRNA synthesis following bortezomib treatment compared to non-induced controls (Figures S6B-C). Additionally, to assess global translation after bortezomib treatment, we performed an HPG assay. We observed marked reduction in HPG signal intensity following BTZ treatment in SLFN11 KMS-27 cells. This result indicates that SLFN11 nucleolar translocation suppresses global translation after bortezomib treatment. Collectively, these findings demonstrate that bortezomib induces the translocation of SLFN11 to nucleoli not only in MM cell lines but also in osteosarcoma U2OS cells, and that this translocation is associated with suppression of ribosomal RNA synthesis.

### SLFN11 MM knockout cells show enhanced sensitivity to bortezomib

To further explore the functional role of SLFN11 in MM cells, we tested whether SLFN11 affects the sensitivity of MM cells proteasome inhibitors. Cell growth experiments revealed that knocking-out SLFN11 both for the KMS-34 and KMS-27 cell lines, showed significantly enhanced sensitivity to bortezomib (Figure 7A). Dose-response experiments using the ATPlite assay confirmed the increased sensitivity to bortezomib in SLFN11 knockout cells (Figure 7B). Conversely, and as expected (37), SLFN11 knockout MM cell lines showed resistance to camptothecin (Figure 7C) and exatecan, the payload of many Antibody Drug Conjugates (ADCs) in clinical trials and preclinical development (Figure 7D). These results confirm SLFN11’s role as a determinant of sensitivity to DNA-damaging agents in MM cells. They also demonstrate the resistance of MM cells to bortezomib in the absence of SLFN11 expression, consistent with a protective role for SLFN11 against proteotoxic stress (9,22).

**Figure 7.**
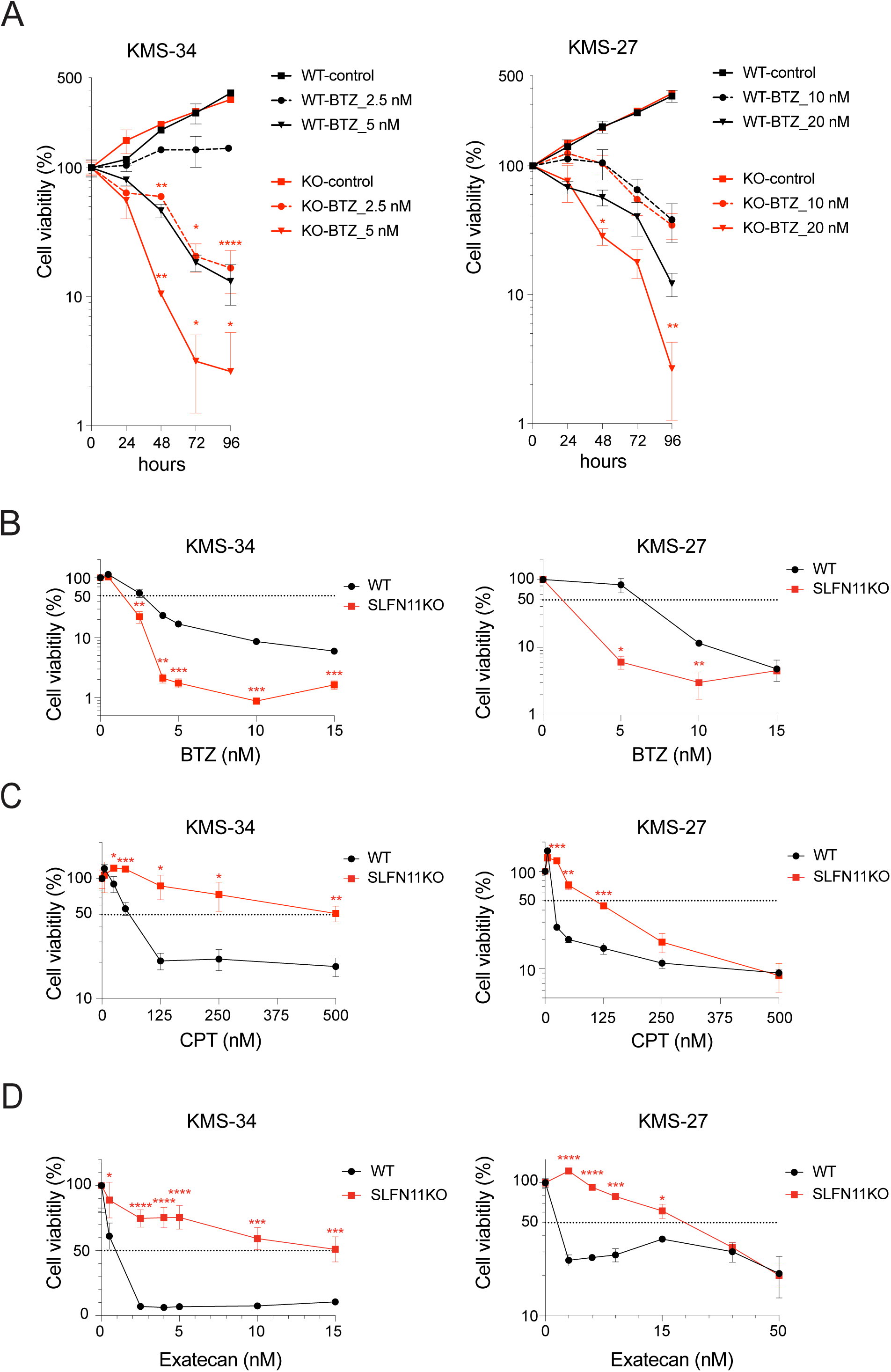
SLFN11 confers resistance to bortezomib and sensitizes to topoisomerase I poisons. (A) Time-course analysis of cell viability in wild-type (WT) and SLFN11 knockout (KO) cells treated with bortezomib (BTZ). Left: KMS-34 cells treated with 2.5 nM or 5 nM BTZ. Right: KMS-27 cells treated with 10 nM or 20 nM BTZ. Cell viability was assessed by trypan blue exclusion at 0, 24, 48, 72, and 96 hours. (B-D) Dose-response curves of WT and SLFN11 KO cells treated for 72 hours with BTZ (0-3 nM) (B), camptothecin (CPT) (0-100 nM) (C) or exatecan (0-3 nM for KMS-34, 0-10 nM for KMS-27) (D). Cell viability was measured using ATP assay. All experiments were performed in triplicate. Data are mean ± SD. *P < 0.05, ** < 0.01, *** < 0.001, **** < 0.0001.

### SLFN11 expression is associated with bortezomib activity in a clinical dataset

To explore the clinical relevance of our experimental findings, we searched published gene expression datasets from randomized clinical trials comparing first-line proteasome inhibitor-containing versus proteasome inhibitor-free therapies in multiple myeloma. The HOVON-65/GMMG-HD4 phase III trial (GSE19784) met these criteria, comparing PAD (bortezomib, doxorubicin, dexamethasone) versus VAD (vincristine, doxorubicin, dexamethasone) induction, followed by high-dose melphalan with autologous stem cell transplantation and maintenance therapy (bortezomib versus thalidomide) in newly diagnosed multiple myeloma with pretreatment gene expression data available for 327 patients (PAD, n=169; VAD, n=158) (40,41). Both regimens share doxorubicin and dexamethasone, differing only in the substitution of vincristine with bortezomib, providing a comparison for bortezomib-specific effects, though a contribution of doxorubicin cannot be fully excluded. *SLFN11* expression was determined using probe 226743_at (Affymetrix HG-U133 Plus 2.0, GPL570) and patients were dichotomized at the cohort median (log₂ = 8.12).

In the VAD arm, low *SLFN11* expression appears a significant adverse prognostic factor. *SLFN11*-low patients showed markedly inferior EFS compared to *SLFN11*-high patients (HR=1.94, 95% CI 1.31–2.88, log-rank P=0.0010; Figure 8A). This adverse prognostic effect was abrogated in the PAD arm (log-rank P=0.11; Figure 8B). Consistent with the pre-specified primary analysis of the original HOVON-65/GMMG-HD4 trial (42), which employed ISS stage-adjusted Cox regression as its primary statistical approach, we applied the same methodology to this subgroup analysis. ISS stage-adjusted Cox regression (n=305, excluding 23 patients with unknown ISS stage). This analysis demonstrated that PAD significantly improved EFS in *SLFN11*-low patients (HR=0.65, 95% CI 0.44–0.96, P=0.030) but not in *SLFN11*-high patients (HR=1.05, 95% CI 0.69–1.60, P=0.809; Figure 8C and S7). Although the formal interaction test did not reach statistical significance (interaction P=0.071), likely reflecting limited statistical power, the directional consistency across all analyses supports the hypothesis that low SLFN11 expression confers selective sensitivity to bortezomib-based therapy.

**Figure 8.**
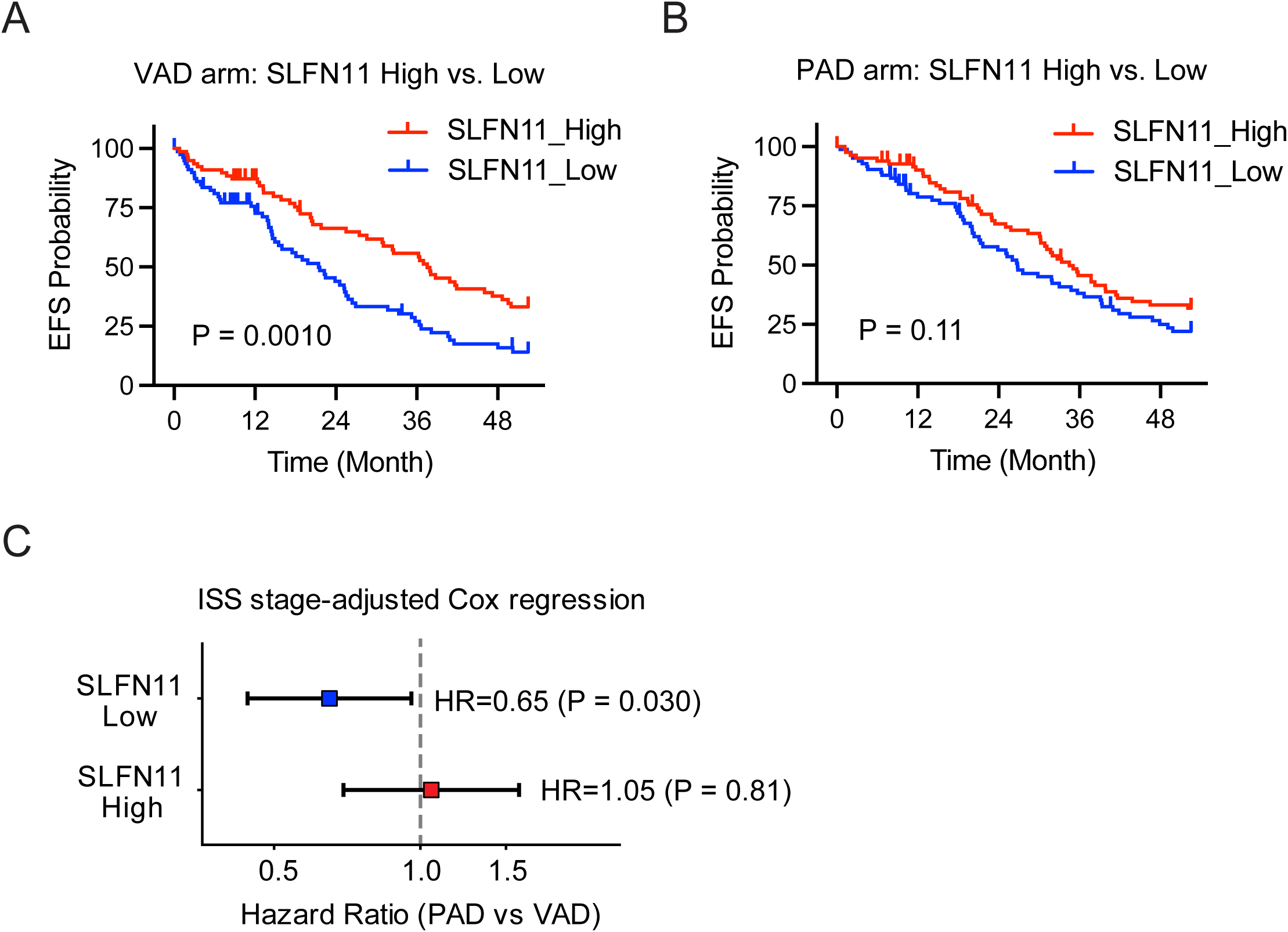
Prediction of the efficacy of bortezomib-based therapy based on *SLFN11* expression in the HOVON-65/GMMG-HD4 clinical trial. Event-free survival was analyzed in 327 newly diagnosed multiple myeloma patients enrolled in the phase III HOVON-65/GMMG-HD4 trial with available pretreatment gene expression data (PAD, n=169; VAD, n=158) (40,41). Patients were dichotomized into *SLFN11*-high and *SLFN11*-low groups at the cohort median (probe 226743_at, log₂ = 8.12). (A, B) Kaplan-Meier analyses with unadjusted log-rank tests; (C) ISS stage-adjusted Cox regression, excluding patients with unknown ISS stage (n=23). (A) EFS in the VAD arm (without bortezomib). (B) EFS in the PAD arm (with bortezomib). (C) ISS stage-adjusted hazard ratios for PAD versus VAD by *SLFN11* group. Survival data were obtained from the GESTURE repository (42). EFS, event-free survival; PAD, bortezomib/doxorubicin/dexamethasone; VAD, vincristine/doxorubicin/dexamethasone; HR, hazard ratio; ISS, International Staging System.

## Discussion

Multiple myeloma (MM) remains an incurable malignancy despite advances in proteasome inhibitor-based therapies. Our study provides the first comprehensive characterization of SLFN11 expression in MM, proposing a previously unrecognized function of SLFN11 in proteotoxic stress response. We demonstrate that *SLFN11* is consistently highly expressed across MM disease stages — intrinsically linked to the plasma cell differentiation program through super-enhancer-mediated transcriptional regulation. We demonstrate that bortezomib induces SLFN11 translocation to nucleoli, where it suppresses ribosomal RNA synthesis. We also show that SLFN11 knockout cells are hypersensitive to bortezomib while being, as expected, resistant to topoisomerase I poisons. This establishes SLFN11 as a dual-function regulator of replication stress and proteotoxic stress responses. These findings reframe SLFN11 not merely as a predictive biomarker for chemotherapies targeting DNA replication, but as an active mediator of proteotoxic stress adaptation in MM, with potential relevance to proteasome inhibitor resistance mechanisms.

MM exhibits remarkably high *SLFN11* expression, comparable to or exceeding AML (18), Ewing sarcoma (20) or mesothelioma (19,43). Notably, this high *SLFN11* expression is maintained throughout malignant transformation and disease progression, suggesting SLN11’s intrinsic relevance for plasma cell biology. This constitutive expression is coordinated with plasma cell-specific transcriptional programs, as *ZBTB38* and *IRF2*, both identified as essential genes regulated by myeloma-specific super-enhancers (35,44), showed strong correlation with *SLFN11* expression. Consistently, blocking super-enhancers with JQ1 and THZ1 suppressed *SLFN11* expression (Figure 3D), providing functional validation of super-enhancer-mediated *SLFN11* regulation in MM.

The constitutive SLFN11 expression in MM is closely linked to plasma cell identity, as evidenced by its strong correlations with plasma cell differentiation markers including SDC1, CD38, and TNFRSF17 (Figure 3A), and its transcriptional regulation by plasma cell-specific super-enhancers (Figure 3D). Notably, SLFN11 expression varies across MM molecular subtypes, with relatively low expression in the CD1 and MF subtypes (Figure 1D). We speculate that in CCND1-driven MM, the balance shifts away from stress-adaptive programs such as SLFN11-mediated translation regulation toward cell cycle progression.

Plasma cells are characterized by exceptionally high immunoglobulin secretory activity, imposing a substantial burden on the protein quality control machinery. The super-enhancer-mediated upregulation of SLFN11 in this context may therefore reflect an adaptive response to this inherently elevated proteotoxic stress, consistent with the enrichment of ER stress response and UPR pathways in *SLFN11*-high MM samples (Figure 4A). Pathway analyses revealed biological signatures relevant to plasma cell biology. *SLFN11*-high samples show enrichment in ER stress response, UPR and ERAD pathways critical for managing high secretory burden. Additionally, *SLFN11*-high samples exhibit reduced ribosomal protein gene expression. Together with our functional data showing SLFN11-mediated suppression of ribosomal RNA synthesis upon bortezomib treatment, this suggests SLFN11 regulates ribosome biogenesis and translation. By tuning down both ribosomal RNA and ribosomal protein synthesis, SLFN11 may function as a regulator of translation to limit cellular stress. These functions may be particularly advantageous in plasma cells, which must reconcile high immunoglobulin production with cellular homeostasis.

Our analyses of proliferative activity show that MM exhibits relatively low *MKI67* expression compared to other cancer types. Both *MKI67* transcript levels and Ki-67 protein expression (PCPI) increase significantly at relapse (Figure 1B, 2C), and the preservation of high *SLFN11* expression combined with increased proliferation in recurrent disease is particularly relevant. Both factors are associated with enhanced sensitivity to DNA-damaging agents (see Figure 7), suggesting that relapsed MM may be more susceptible to DNA-damaging therapies, such as TOP1-ADCs (with exatecan and deruxtecan payloads) than newly diagnosed disease.

In untreated conditions, SLFN11 is loosely bound to chromatin (detergent soluble) throughout the nucleus and largely excluded from nucleoli (14,19,39,45). Upon replicative DNA damage by chemotherapeutic agents, SLFN11 forms nuclear foci in replication factories (7,29,39,46). By contrast, bortezomib induces SLFN11 translocation to the fibrillar center of nucleoli. This nucleolar binding appears functionally significant. Following bortezomib treatment, we find that SLFN11 suppresses ribosomal RNA synthesis. The ability of SLFN11 to down-regulate rDNA synthesis aligns with a recent publication demonstrating that SLFN11 can bind ribosomal DNA and inhibit rRNA synthesis (9). Our results extend these observations to proteotoxic stress induced by proteasome inhibition, suggesting that SLFN11-mediated rRNA synthesis suppression may represent a conserved mechanism that operates across different types of cellular stresses. Thus, we propose a dual-function model for SLFN11 in multiple myeloma. Under DNA damage stress induced by TOP1 poisons (or chemotherapy-induced replication stress), SLFN11 binds to and blocks replication forks, leading to the death of cells with irreparable damage. In contrast, under proteotoxic stress induced by proteasome inhibition, SLFN11 translocates to nucleoli and suppresses ribosomal RNA synthesis, thereby reducing overall protein synthesis (see Figure S6) and mitigating the accumulation of misfolded proteins.

This model has therapeutic implications for multiple myeloma treatment. *SLFN11*-high myeloma cells may possess an intrinsic protective mechanism against proteasome inhibition through SLFN11-mediated control of ribosomal RNA synthesis, potentially conferring relative resistance to bortezomib. In contrast, *SLFN11*-low myeloma cells lack this protective mechanism and continue ribosomal RNA synthesis despite proteasome inhibition, potentially leading to enhanced sensitivity to bortezomib. This hypothesis is consistent with our observation that *SLFN11*-low myeloma samples show upregulation of ubiquitin-proteasome system components, suggesting a greater dependency on proteasome function for survival. Based on our model, *SLFN11*-low patients might benefit most from bortezomib-based regimens. For patients with very high *SLFN11* expression, combining proteasome inhibitors with agents targeting ribosome biogenesis might further abolish rRNA and protein synthesis. Additionally, given SLFN11’s established role in sensitizing cancer cells to DNA-damaging agents (7,29), *SLFN11*-high myeloma patients might benefit from DNA-damaging agents targeting S-phase cells in their treatment regimens. We confirmed this concept functionally by demonstrating that *SLFN11* knockout in MM cell lines (KMS-27 and KMS-34) conferred resistance to topoisomerase I poisons (camptothecin and exatecan), directly validating SLFN11’s role as a determinant of DNA-damaging agent sensitivity in MM cells (Figure 7C, 7D). This rationale is further strengthened by our observation that proliferative activity (PCPI) increases during disease progression while *SLFN11* expression remains high, potentially enhancing susceptibility to DNA-damaging therapies in the relapsed setting. The strong correlation between *SLFN11* and *BCMA* expression (r = 0.37) provides additional rationale for BCMA-targeted therapies in *SLFN11*-high patients. Topoisomerase I (TOP1) poisons are selectively toxic to replicating cells and are increasingly used as payloads for ADCs (47,48). While current anti-BCMA ADCs utilize microtubule inhibitor payloads (49,50), TOP1 payloads such as deruxtecan and exatecan could offer superior efficacy in *SLFN11*-high MM patients, particularly at relapse where both *SLFN11* expression and proliferative activity are high.

Consistent with this mechanistic model, a retrospective subgroup analysis of the HOVON-65/GMMG-HD4 trial provides hypothesis-generating clinical evidence. In the VAD arm (without bortezomib), low *SLFN11* expression was a significant adverse prognostic factor (P=0.0010), yet this effect was abrogated in the PAD arm (P=0.11). ISS stage-adjusted Cox regression further demonstrated selective EFS benefit from bortezomib specifically in *SLFN11*-low patients (HR=0.65, P=0.030), supporting the interpretation that *SLFN11*-low MM cells, lacking the protective rRNA synthesis suppression mechanism, are selectively sensitized to bortezomib-induced proteotoxic stress. Prospective validation in larger cohorts with standardized SLFN11 measurement is required.

While previous studies established PCPI as a prognostic factor using cross-sectional analyses, our sequential assessment revealed that PCPI progressively increases with each relapse—a pattern not systematically documented previously (33,51). Combined with maintained *SLFN11* expression, this suggests a therapeutic window in relapsed disease where both factors converge to enhance DNA-damaging agent susceptibility.

Several limitations of our study should be acknowledged. Although we demonstrate SLFN11-mediated suppression of ribosomal RNA synthesis following bortezomib treatment, the detailed molecular mechanisms underlying this function remain to be elucidated. Our mechanistic studies were conducted in cell line models, and confirmation in primary patient samples are needed to strengthen our conclusions, particularly given the difference between high *SLFN11* expression in MM cell lines (∼33%) and patient-derived samples (97.2%). Additionally, comprehensive evaluation of SLFN11’s role in MM progression and its predictive value for proteasome inhibitor response require further investigations, including prospective clinical studies. In vivo studies using SLFN11 knockout MM models are warranted to further define its role in disease progression.

## Supporting information

Supplementary Methods, Figure Legends, and References

Figure S1. SLFN11 expression

Figure S2. SLFN11/CD138 patterns

Figure S3. SLFN11 correlations

Figure S4. Pathways and GSEA

Figure S5. SLFN11 in U2OS

Figure S6. rRNA and translation

Figure S7. HOVON-65 EFS

## Acknowledgements

This work was supported by JSPS KAKENHI Grant-in-Aid for Scientific Research (C) [25K11676] to Y.A. Our studies are supported by the intramural program of the US National Cancer Institute, NIH, Bethesda, Maryland (Z01-BC-006150). SLFN11-targeting CRISPR constructs were kindly provided by Dr. Junko Murai.

## Data Availability Statement

Public datasets analyzed in this study are available through UCSC Xena Functional Genomics Explorer (https://xenabrowser.net) for TCGA and MMRF CoMMpass Study data, and Gene Expression Omnibus (GEO) for microarray datasets (GSE5900 and GSE2658). Additional raw data are available from the corresponding author upon reasonable request.

## References

1. Cowan AJ, Green DJ, Kwok M, Lee S, Coffey DG, Holmberg LA, et al. Diagnosis and Management of Multiple Myeloma: A Review. JAMA 2022;327:464–77

2. Zhan F, Huang Y, Colla S, Stewart JP, Hanamura I, Gupta S, et al. The molecular classification of multiple myeloma. Blood 2006;108:2020–8

3. Maura F, Rajanna AR, Ziccheddu B, Poos AM, Derkach A, Maclachlan K, et al. Genomic Classification and Individualized Prognosis in Multiple Myeloma. J Clin Oncol 2024;42:1229–40

4. Skerget S, Penaherrera D, Chari A, Jagannath S, Siegel DS, Vij R, et al. Comprehensive molecular profiling of multiple myeloma identifies refined copy number and expression subtypes. Nat Genet 2024;56:1878–89

5. Soekojo CY, Chng WJ. Treatment horizon in multiple myeloma. Eur J Haematol 2022;109:425–40

6. Jo U, Pommier Y. Structural, molecular, and functional insights into Schlafen proteins. Experimental & Molecular Medicine 2022

7. Murai J, Thomas A, Miettinen M, Pommier Y. Schlafen 11 (SLFN11), a restriction factor for replicative stress induced by DNA-targeting anti-cancer therapies. Pharmacol Ther 2019;201:94–102

8. Boon NJ, Oliveira RA, Korner PR, Kochavi A, Mertens S, Malka Y, et al. DNA damage induces p53-independent apoptosis through ribosome stalling. Science 2024;384:785–92

9. Ogawa A, Izumikawa K, Tate S, Isoyama S, Mori M, Fujiwara K, et al. SLFN11-mediated ribosome biogenesis impairment induces TP53-independent apoptosis. Mol Cell 2025;85:894–912 e10

10. Allison Stewart C, Tong P, Cardnell RJ, Sen T, Li L, Gay CM, et al. Dynamic variations in epithelial-to-mesenchymal transition (EMT), ATM, and SLFN11 govern response to PARP inhibitors and cisplatin in small cell lung cancer. Oncotarget 2017;8:28575–87

11. Murai J, Ceribelli M, Fu H, Redon CE, Jo U, Murai Y, et al. Schlafen 11 (SLFN11) Kills Cancer Cells Undergoing Unscheduled Re-replication. Mol Cancer Ther 2023;22:985–95

12. Murai J, Feng Y, Yu GK, Ru Y, Tang SW, Shen Y, et al. Resistance to PARP inhibitors by SLFN11 inactivation can be overcome by ATR inhibition. Oncotarget 2016;7:76534–50

13. Takashima T, Taniyama D, Sakamoto N, Yasumoto M, Asai R, Hattori T, et al. Schlafen 11 predicts response to platinum-based chemotherapy in gastric cancers. Br J Cancer 2021;125:65–77

14. Taniyama D, Sakamoto N, Takashima T, Takeda M, Pham QT, Ukai S, et al. Prognostic impact of Schlafen 11 in bladder cancer patients treated with platinum-based chemotherapy. Cancer Sci 2022;113:784–95

15. Willis SE, Winkler C, Roudier MP, Baird T, Marco-Casanova P, Jones EV, et al. Retrospective analysis of Schlafen11 (SLFN11) to predict the outcomes to therapies affecting the DNA damage response. Br J Cancer 2021;125:1666–76

16. Zhang B, Ramkumar K, Cardnell RJ, Gay CM, Stewart CA, Wang WL, et al. A wake-up call for cancer DNA damage: the role of Schlafen 11 (SLFN11) across multiple cancers. Br J Cancer 2021;125:1333–40

17. Zoppoli G, Regairaz M, Leo E, Reinhold WC, Varma S, Ballestrero A, et al. Putative DNA/RNA helicase Schlafen-11 (SLFN11) sensitizes cancer cells to DNA-damaging agents. Proceedings of the National Academy of Sciences of the United States of America 2012;109:15030–5

18. Murai Y, Jo U, Murai J, Fukuda S, Takebe N, Pommier Y. Schlafen 11 expression in human acute leukemia cells with gain-of-function mutations in the interferon-JAK signaling pathway. iScience 2021;24:103173

19. Kaczorowski M, Ylaya K, Chlopek M, Taniyama D, Pommier Y, Lasota J, et al. Immunohistochemical Evaluation of Schlafen 11 (SLFN11) Expression in Cancer in the Search of Biomarker-Informed Treatment Targets: A Study of 127 Entities Represented by 6658 Tumors. Am J Surg Pathol 2024

20. Tang SW, Bilke S, Cao L, Murai J, Sousa FG, Yamade M, et al. SLFN11 Is a Transcriptional Target of EWS-FLI1 and a Determinant of Drug Response in Ewing Sarcoma. Clin Cancer Res 2015;21:4184–93

21. Li M, Kao E, Gao X, Sandig H, Limmer K, Pavon-Eternod M, et al. Codon-usage-based inhibition of HIV protein synthesis by human schlafen 11. Nature 2012;491:125–8

22. Murai Y, Jo U, Murai J, Jenkins LM, Huang SN, Chakka S, et al. SLFN11 Inactivation Induces Proteotoxic Stress and Sensitizes Cancer Cells to Ubiquitin Activating Enzyme Inhibitor TAK-243. Cancer Res 2021;81:3067–78

23. Otsuki T, Yata K, Takata-Tomokuni A, Hyodoh F, Miura Y, Sakaguchi H, et al. Expression of protein gene product 9.5 (PGP9.5)/ubiquitin-C-terminal hydrolase 1 (UCHL-1) in human myeloma cells. Br J Haematol 2004;127:292–8

24. Kim JM, Kee Y, Gurtan A, D’Andrea AD. Cell cycle-dependent chromatin loading of the Fanconi anemia core complex by FANCM/FAAP24. Blood 2008;111:5215–22

25. Caicedo HH, Hashimoto DA, Caicedo JC, Pentland A, Pisano GP. Overcoming barriers to early disease intervention. Nat Biotechnol 2020;38:669–73

26. Goldman MJ, Craft B, Hastie M, Repecka K, McDade F, Kamath A, et al. Visualizing and interpreting cancer genomics data via the Xena platform. Nat Biotechnol 2020;38:675–8

27. Lachmann A, Xie Z, Ma’ayan A. blitzGSEA: efficient computation of gene set enrichment analysis through gamma distribution approximation. Bioinformatics 2022;38:2356–7

28. Ding L, Bailey MH, Porta-Pardo E, Thorsson V, Colaprico A, Bertrand D, et al. Perspective on Oncogenic Processes at the End of the Beginning of Cancer Genomics. Cell 2018;173:305–20 e10

29. Jo U, Murai Y, Takebe N, Thomas A, Pommier Y. Precision Oncology with Drugs Targeting the Replication Stress, ATR, and Schlafen 11. Cancers (Basel) 2021;13

30. Yerushalmi R, Woods R, Ravdin PM, Hayes MM, Gelmon KA. Ki67 in breast cancer: prognostic and predictive potential. Lancet Oncol 2010;11:174–83

31. Gastinne T, Leleu X, Duhamel A, Moreau AS, Franck G, Andrieux J, et al. Plasma cell growth fraction using Ki-67 antigen expression identifies a subgroup of multiple myeloma patients displaying short survival within the ISS stage I. Eur J Haematol 2007;79:297–304

32. Zhan F, Barlogie B, Arzoumanian V, Huang Y, Williams DR, Hollmig K, et al. Gene-expression signature of benign monoclonal gammopathy evident in multiple myeloma is linked to good prognosis. Blood 2007;109:1692–700

33. Ely S, Forsberg P, Ouansafi I, Rossi A, Modin A, Pearse R, et al. Cellular Proliferation by Multiplex Immunohistochemistry Identifies High-Risk Multiple Myeloma in Newly Diagnosed, Treatment-Naive Patients. Clin Lymphoma Myeloma Leuk 2017;17:825–33

34. Kassambara A, Herviou L, Ovejero S, Jourdan M, Thibaut C, Vikova V, et al. RNA-sequencing data-driven dissection of human plasma cell differentiation reveals new potential transcription regulators. Leukemia 2021;35:1451–62

35. Jia Y, Zhou J, Tan TK, Chung TH, Wong RWJ, Chooi JY, et al. Myeloma-specific superenhancers affect genes of biological and clinical relevance in myeloma. Blood Cancer J 2021;11:32

36. Xiong S, Zhou J, Tan TK, Chung TH, Tan TZ, Toh SH, et al. Super enhancer acquisition drives expression of oncogenic PPP1R15B that regulates protein homeostasis in multiple myeloma. Nat Commun 2024;15:6810

37. Elloumi F, Tlemsani C, Heske CM, Pongor L, Khandagale P, Varma S, et al. Protocol to set up the CellMinerCDB pharmacogenomics analysis web application with custom cell line data. STAR Protoc 2025;6:104076

38. Tlemsani C, Heske CM, Elloumi F, Pongor L, Khandagale P, Varma S, et al. Sarcoma_CellminerCDB: A tool to interrogate the genomic and functional characteristics of a comprehensive collection of sarcoma cell lines. iScience 2024;27:109781

39. Murai J, Tang SW, Leo E, Baechler SA, Redon CE, Zhang H, et al. SLFN11 Blocks Stressed Replication Forks Independently of ATR. Mol Cell 2018;69:371–84 e6

40. Broyl A, Hose D, Lokhorst H, de Knegt Y, Peeters J, Jauch A, et al. Gene expression profiling for molecular classification of multiple myeloma in newly diagnosed patients. Blood 2010;116:2543–53

41. Ubels J, Sonneveld P, van Beers EH, Broijl A, van Vliet MH, de Ridder J. Predicting treatment benefit in multiple myeloma through simulation of alternative treatment effects. Nat Commun 2018;9:2943

42. Sonneveld P, Schmidt-Wolf IG, van der Holt B, El Jarari L, Bertsch U, Salwender H, et al. Bortezomib induction and maintenance treatment in patients with newly diagnosed multiple myeloma: results of the randomized phase III HOVON-65/ GMMG-HD4 trial. J Clin Oncol 2012;30:2946–55

43. Rathkey D, Khanal M, Murai J, Zhang J, Sengupta M, Jiang Q, et al. Sensitivity of Mesothelioma Cells to PARP Inhibitors Is Not Dependent on BAP1 but Is Enhanced by Temozolomide in Cells With High-Schlafen 11 and Low-O6-methylguanine-DNA Methyltransferase Expression. J Thorac Oncol 2020;15:843–59

44. de Matos Simoes R, Shirasaki R, Downey-Kopyscinski SL, Matthews GM, Barwick BG, Gupta VA, et al. Genome-scale functional genomics identify genes preferentially essential for multiple myeloma cells compared to other neoplasias. Nat Cancer 2023;4:754–73

45. Coleman N, Zhang B, Byers LA, Yap TA. The role of Schlafen 11 (SLFN11) as a predictive biomarker for targeting the DNA damage response. Br J Cancer 2021;124:857–9

46. Mu Y, Lou J, Srivastava M, Zhao B, Feng XH, Liu T, et al. SLFN11 inhibits checkpoint maintenance and homologous recombination repair. EMBO Rep 2016;17:94–109

47. Jiang Y, Xu Y, He J, Sui L, Li T, Xia N, et al. Uncovering potential targets for antibody-drug conjugates in the treatment of gynecologic malignancies. Front Pharmacol 2025;16:1525733

48. Thomas A, Pommier Y. Targeting Topoisomerase I in the Era of Precision Medicine. Clin Cancer Res 2019;25:6581–9

49. Dimopoulos MA, Beksac M, Pour L, Delimpasi S, Vorobyev V, Quach H, et al. Belantamab Mafodotin, Pomalidomide, and Dexamethasone in Multiple Myeloma. N Engl J Med 2024;391:408–21

50. Trudel S, Stewart AK. A Belantamab Mafodotin Revival in Multiple Myeloma Therapy. N Engl J Med 2024;391:461–2

51. Aljama MA, Sidiqi MH, Lakshman A, Dispenzieri A, Jevremovic D, Gertz MA, et al. Plasma cell proliferative index is an independent predictor of progression in smoldering multiple myeloma. Blood Adv 2018;2:3149–54

