## Supplementary Methods, Figure Legends, and References for "Schlafen 11 (SLFN11) overexpression in multiple myeloma and nucleolar translocation in response to bortezomib"

**Supplemental Methods**

**Cell Lines and Culture Conditions**

KMS-27 (RRID:CVCL_2993) and KMS-34 (RRID:CVCL_2996) multiple myeloma cell lines (1) were obtained directly from the Japanese Collection of Research Bioresources Cell Bank and maintained in RPMI-1640 medium (Gibco #11875093) supplemented with 10% fetal bovine serum (FBS) (Gibco #A5256701), penicillin/streptomycin, and 2.5 μg/ml amphotericin B (FUJIFILM #019-23891). MM.1S multiple myeloma cell line (RRID:CVCL_8792), NCI-H295R adrenocortical carcinoma cell line (RRID:CVCL_0458), and U2OS osteosarcoma cell line (RRID:CVCL_0042) were obtained directly from the American Type Culture Collection. These cell lines were used within 6 months of receipt. MM.1S cells were cultured in the same medium as KMS cells. U2OS cells were maintained in McCoy's 5a medium (Gibco #16600082) supplemented with 10% FBS and penicillin/streptomycin. NCI-H295R cells were cultured in DMEM/F12 medium (Gibco #11320033) supplemented with 2.5% Nu-Serum (Corning #355100) and ITS+ Premix (Corning #354352). All cell lines were regularly tested for mycoplasma contamination using the MycoAlert Mycoplasma Detection Kit (Lonza #LT07-318) and maintained at 37°C in a humidified atmosphere with 5% CO₂.

**Patient Specimens**

Bone marrow aspirates and biopsies from multiple myeloma patients were obtained with approval from the Ethics Committee of the Jikei University School of Medicine (approval number 37-008). Inclusion criteria: Patients diagnosed with multiple myeloma according to International Myeloma Working Group criteria with adequate bone marrow specimens for analysis. Exclusion criteria: Patients with concurrent other hematological malignancies or insufficient sample quality for immunohistochemical analysis. For this retrospective study, informed consent was obtained directly from patients when possible. An opt-out approach was applied for deceased patients or those lost to follow-up, in accordance with institutional ethical guidelines approved specifically for retrospective analyses of archived clinical specimens. Normal bone marrow tissue arrays (BN29011a and BM241b) were purchased from TissueArray.Com (Derwood, MD, USA). Sections of 2-4 μm thickness were prepared from formalin-fixed paraffin-embedded specimens for immunohistochemical analysis. Sequential bone marrow samples were collected from patients at initial diagnosis and during disease progression or relapse following treatment. Normal bone marrow tissue arrays were used as controls.

**Dual Immunohistochemical Staining for CD138 and Ki-67**

To evaluate the expression patterns of SLFN11 and CD138 in patient bone marrow samples, dual immunohistochemical staining was performed. Deparaffinized FFPE sections underwent antigen retrieval in high pH buffer (DAKO #K8004) at 95°C for 20 minutes. After blocking endogenous peroxidase activity with 0.3% H₂O₂, sections were incubated with anti-SLFN11 antibody (Santa Cruz, #sc-515071, 1:100 dilution) at room temperature for 60 minutes, followed by visualization using a polymer detection system (Histofine Simple Stain MAX-PO, Nichirei Biosciences #414341) and DAB chromogen (brown). Subsequently, sections were incubated with anti-CD138 antibody (Proteintech, 10593-1-AP, 1:1000 dilution) at 4°C overnight, followed by Goat Anti-Rabbit IgG antibody, VECTASTAIN ABC-AP Kit (Vector Laboratories #AK-5000), and Vector Blue chromogen (blue). Sections were counterstained with nuclear fast red, dehydrated, and mounted for microscopic analysis. Stained slides were examined to quantify the proportions of CD138+/SLFN11+, CD138+/SLFN11-, CD138-/SLFN11+, and CD138-/SLFN11- cell populations. Cell populations were quantified by counting 1,000 cells per sample (including both CD138-positive and CD138-negative cells), except for samples with low cell density where 500 cells were counted. For normal bone marrow samples, 200 CD138-positive cells were counted per sample.

For CD138/Ki-67 dual staining to assess proliferative activity, the same protocol was used with anti-Ki-67 antibody (Santa Cruz, #sc-101861, 1:100 dilution) replacing anti-SLFN11 antibody in the first staining step. The plasma cell proliferation index (PCPI) was calculated as the percentage of CD138-positive cells co-expressing Ki-67, using the same cell counting criteria described above.

**Immunofluorescence analysis with pre-extraction**

For multiple myeloma cell lines, cells were applied to Superfrost Plus microscope slides (Electron Microscopy Sciences, 71869-10) using a Cytospin at 800 rpm for 4 minutes. To detect the chromatin-bound proteins (For Figures 5D and 5E), the attached cells were briefly pretreated with ice-cold 0.1% Triton X-100 in PBS for 1 minute, followed by fixation in 4% paraformaldehyde in PBS for 10 minutes at room temperature (2). For U2OS cell line, cells (5×10^4^ cells) were seeded on 8-well chamber slides (Nunc™ Lab-Tek™ II CC2™, Thermo Fisher, #154941) except for experiments in Figures S5A and S5B, where 2000 cells were seeded per well in 384-well plate (Perkin Elmer CellCarrier 384 plates, #6057300). Cells were treated with drugs as described in the figure legends. Cells were then pre-extracted in 0.1% Triton X-100 cytoskeleton (CSK) buffer (10 mM Tris [pH 6.8], 100 mM NaCl, 300 mM sucrose, MgCl2, 1 mM EGTA, 1 mM EDTA, and 0.1% Triton X-100) for 5 min on ice, following fixation in 4% PFA for 15 min (3). For Figures 6A and 6C, the attached cells were fixed with 4% paraformaldehyde for 10 min followed by permeabilization with 0.1% Triton X-100/PBS for 15 min. After fixation, cells were blocked with 4% bovine serum albumin (BSA) in PBST for 10 minutes, then incubated overnight at 4 °C in a humidified chamber with primary antibodies diluted 1:1000 in 4% BSA/PBST. The primary antibodies used were anti-SLFN11 (Santa Cruz Biotechnology, D2) and anti-Nucleolin (Abcam, EPR7952). Following PBST washes, cells were incubated with Alexa Fluor 488-conjugated goat anti-mouse IgG or Alexa Fluor 568-conjugated goat anti-rabbit IgG (both 1:1000 in 5% BSA/PBST) for 2 hours at room temperature. After additional PBST washes, the cells were mounted using Vectashield containing DAPI (VECTOR, H-1200). Imaging was performed using a Nikon SoRa spinning disk confocal microscope equipped with a 60× objective lens. For the experiments in Figures 5F, super-resolution imaging was performed, and post-acquisition deconvolution was applied exclusively to these images using NIS-Elements software (Nikon) to further enhance resolution and contrast.

**Immunofluorescence Microscopy Image Analysis**

Fiji (ImageJ, NIH ; RRID:SCR_002285) software was used to quantify signal intensities at the single-cell level. Nuclear regions were identified by DAPI staining and used to define regions of interest across isogenic cell lines for the extraction of mean signal intensities. These values were used to generate individual plots. The processed data were subsequently imported into GraphPad Prism 11 (GraphPad Software; RRID:SCR_002798) for graphical representation.

**Automated image acquisition and image analysis**

In Figure S5A and S5B, Stained 384-well plates were imaged on a Yokogawa CV7000S high-throughput spinning-disk confocal microscope using a 40× air objective (NA 0.95). DAPI and chromatin-bound SLFN11 were detected with 405 nm and 561 nm excitation, respectively. Single-plane images were acquired sequentially with a 16-bit sCMOS camera (2×2 binning; pixel size, 0.325 μm) using appropriate bandpass filters. Background and illumination corrections were performed using Yokogawa software, and images were exported as TIFF files. Image analysis was performed in Columbus 2.7.1 (PerkinElmer). Nuclei were segmented based on DAPI signal, and nuclear SLFN11 fluorescence intensity was quantified after treatment with camptothecin (CPT) or bortezomib (BTZ) for 4 hours. Nuclei with roundness <0.775 or touching the image border were excluded. Mean nuclear SLFN11 intensity was calculated per well and exported as text files for downstream analysis. This workflow was used in Figures S5A and S5B.

**Western Blotting**

For protein expression analysis, cells were lysed in RIPA buffer (50 mM Tris-HCl pH 7.4, 150 mM NaCl, 1% NP-40, 0.5% sodium deoxycholate, 0.1% SDS, 1 mM EDTA) supplemented with protease inhibitor cocktail (Roche #11836170001). Protein concentrations were determined using the Bio-Rad Protein Assay Kit (Bio-Rad #5000001). Equal amounts of protein (10 μg) were resolved by 5 to 20% SDS-PAGE and transferred to PVDF membranes (Millipore #IPVH00005). After blocking with 5% non-fat dry milk in TBS-T, membranes were incubated with primary antibodies against SLFN11 (Santa Cruz #sc-515071, 1:1000) or β-actin (Sigma-Aldrich #A551, 1:5000) overnight at 4°C. Following incubation with HRP-conjugated secondary antibodies (Santa Cruz #sc-2357, #sc-2004), immunoreactive bands were visualized using ECL detection reagent (Atto #WSE-7110) and analyzed using Fiji software (NIH).

**Treatment with Super Enhancer Inhibitors**

Multiple myeloma cell lines were treated with JQ1 (BET inhibitor, Sigma-Aldrich #SML1524-5MG) or THZ1·2HCl (CDK7 inhibitor, Selleck #S7549) to evaluate their effects on SLFN11, ZBTB38, and IRF2 protein expression. Stock solutions were prepared in DMSO. KMS-34 cells were treated with JQ1 (500 nM) or THZ1 (200 nM), and KMS-27 cells were treated with JQ1 (1 μM) or THZ1 (50 nM) for 0, 12, 24, and 48 hours. Cells were harvested at each time point for Western blot analysis as described above.

**CRISPR-Cas9-mediated SLFN11 Knockout**

SLFN11 knockout (KO) clones were generated in KMS-34 cells using the CRISPR-Cas9 system. SLFN11-targeting CRISPR constructs were kindly provided by Dr. Junko Murai (4) and modified according to the published protocol with minor adaptations. Briefly, KMS-27and KMS-34 cells were transfected with the CRISPR constructs using Nucleofector II Device with Nucleofector Kit V (Amaxa #VCA-1003) and program X-005 (5). Following transfection, cells were cultured in drug-free medium for 48 hours and subsequently selected with 0.25 μg/ml puromycin (Thermo Fisher Scientific #A1113803) for one week. Single-cell clones were isolated by limiting dilution, expanded, and screened for SLFN11 knockout by Western blotting and immunofluorescence analysis.

**Cell Viability Assays**

Cell viability was assessed using either manual cell counting with trypan blue exclusion or CellTiter-Glo Luminescent Cell Viability Assay (Promega #G7570). For time-course experiments with BTZ (Figure 7A), wild-type (WT) and SLFN11 knockout (KO) cells were seeded in 24-well plates at a density of 2×10⁵ cells/mL and treated with indicated concentrations of BTZ. At 0, 24, 48, 72, and 96 hours, viable cells were counted after trypan blue staining.

For dose-response experiments (Figure 7B-D), WT and SLFN11 KO cells were seeded in 96-well plates according to CellTiter-Glo protocol recommendations and treated with various concentrations of BTZ (0-3 nM), CPT (Sigma-Aldrich #390238, 0-100 nM), or exatecan (Sigma-Aldrich #SML3519, 0-3 nM for KMS-34 or 0-10 nM for KMS-27) for 72 hours. Cell viability was measured using CellTiter-Glo reagent according to the manufacturer's instructions, and luminescence was recorded using a plate reader. All experiments were performed in triplicate. Data were normalized to untreated control cells (100% viability).

**Doxycycline-inducible SLFN11 Expression System**

U2OS cells stably expressing SLFN11 under the control of a doxycycline-inducible promoter were generated via lentiviral transduction using the pLVX-TetOne-Blasticidin-3XFlag-SLFN11 construct. Forty-eight hours after infection, transduced cells were selected with 10 µg/mL blasticidin (Gibco). Cells were treated with doxycycline (DOX, Thermo Fisher, 5 ug/mL) for 72 hours to induce *SLFN11*.

**Ribosomal RNA Synthesis Assay**

Followed by treating cells with drugs as described in the figure legends, cells were incubated with 5-ethynyl uridine (EU) for 1 h (5 mM for Fig. 6C; 1 mM for Fig. S6), and nascent RNA was detected using the Click-iT RNA Alexa Fluor 488 Imaging Kit (Thermo Fisher, C10329), following the manufacturer’s instructions. EU fluorescence signal intensities were quantified using Fiji. All data were visualized using GraphPad Prism 11 (GraphPad Software).

**Protein synthesis assay**

To evaluate global protein synthesis, the Click-iT™ HPG Alexa Fluor™ 488 Protein Synthesis Assay Kit (Thermo Fisher, # C10329) was used according to the manufacturer's instructions. Cells were treated with 100 nM BTZ for 2 or 4 h. To label nascent proteins without inducing amino acid starvation, 50 μM HPG was added directly to the culture medium during the final hour of BTZ treatment. After fixation and permeabilization, the click reaction was performed to detect incorporated HPG. Fluorescence signals were analyzed using a confocal microscope. HPG fluorescence signal intensities were quantified using Fiji. All data were visualized using GraphPad Prism 11.

**Analysis of Public Gene Expression Datasets**

Public gene expression datasets were analyzed to investigate *SLFN11* expression across cancer types and multiple myeloma subtypes. The MMRF CoMMpass dataset (initial and relapsed bone marrow samples, N=844) and TCGA Pan-Cancer dataset were analyzed and visualized using UCSC Xenabrowser (https://xenabrowser.net, data accessed February 2025; RRID:SCR_018938) (6,7). Pre-processed RNA-seq data with STAR aligner (RRID:SCR_004463) mapping and FPKM-UQ normalization were downloaded and analyzed. Microarray data of normal plasma cells and sequential stages of myeloma progression were downloaded from the Gene Expression Omnibus (GEO ; RRID:SCR_005012) database. Specifically, we utilized datasets GSE5900 and GSE2658 for analyzing *SLFN11* expression across normal plasma cells (NPC), monoclonal gammopathy of undetermined significance (MGUS), smoldering multiple myeloma (SMM), and symptomatic multiple myeloma (MM) (8,9). Correlation analysis between drug sensitivity and *SLFN11* expression from the Cancer Therapeutics Response Portal (CTRP) database was performed and visualized using CellMiner CDB (<https://discover.nci.nih.gov/rsconnect/cellminercdb/>) (10). Gene expression values were normalized as log2(fpkm-uq+1) for RNA-seq data analysis.

**Retrospective Clinical Subgroup Analysis (HOVON-65/GMMG-HD4)**

Gene expression data from the HOVON-65/GMMG-HD4 Phase III trial were obtained from Gene Expression Omnibus (GSE19784) (11). Event-free survival (EFS) data and treatment group assignments for 327 patients for whom gene expression profiling data were available (PAD group: n=169, VAD group: n=158) were obtained from the GESTURE repository (github.com/jubels/GESTURE) (12). SLFN11 expression levels were quantified using probe 226743_at on the Affymetrix HG-U133 Plus 2.0 array (GPL570) and verified using the hgu133plus2.db Bioconductor annotation package in R. Patients were divided into *SLFN11* high-expression and *SLFN11* low-expression groups based on the cohort median (log₂ = 8.12). Kaplan-Meier survival curves were constructed and compared using the log-rank test. Patients with unknown ISS stage (n = 23) were excluded, and an ISS-adjusted Cox proportional hazards regression analysis was performed. The interaction between the SLFN11 group and the treatment group was evaluated using a Cox model that included an SLFN11 group × treatment group interaction term. All analyses were performed using the survival and survminer packages in R version 4.4.3.

**Pathway Analysis and Gene Set Enrichment Analysis**

Differential gene expression (DGE) analysis and Gene Set Enrichment Analysis (GSEA) were performed using the analytical functions available in Xenabrowser (7). Samples from the MMRF CoMMpass study, including both newly diagnosed and relapsed bone marrow specimens, were divided into SLFN11-high and SLFN11-low groups based on median SLFN11 expression. DGE analysis was conducted using Xenabrowser's limma-voom implementation (13,14).

Gene Ontology (GO) enrichment analysis was performed using Xenabrowser's Enrichr integration to identify significantly enriched biological processes, molecular functions, and cellular components in differentially expressed genes between *SLFN11*-high and *SLFN11*-low samples (15,16). Gene Set Enrichment Analysis was conducted using the blitzGSEA function within Xenabrowser with gene sets from the Molecular Signatures Database (MSigDB), including both Hallmark and Ontology gene sets (17). Positive Normalized Enrichment Scores (NES) indicate enrichment in *SLFN11*-high samples, while negative scores indicate enrichment in *SLFN11*-low samples.

Heatmaps of differentially expressed genes were generated using Xenabrowser's visualization tools to display expression patterns across samples, with particular focus on genes related to ER stress response, unfolded protein response, ribosomal proteins, and ubiquitin-proteasome system components.

**Statistical Analyses**

Statistical analyses were performed using GraphPad Prism 11 (GraphPad Software ; RRID:SCR_002798). Comparisons between two groups were conducted using either unpaired t-test or Mann-Whitney U test depending on data distribution. For multiple group comparisons, one-way ANOVA with Tukey's post-hoc test or Kruskal-Wallis test with Dunn's post-hoc test was applied as appropriate. Correlations between gene expression levels were evaluated using Pearson's correlation coefficient. For all analyses, P values less than 0.05 were considered statistically significant. Details regarding data presentation are provided in the respective figure legends.

**Supplemental References**

1.	Otsuki T, Yata K, Takata-Tomokuni A, Hyodoh F, Miura Y, Sakaguchi H*, et al.* Expression of protein gene product 9.5 (PGP9.5)/ubiquitin-C-terminal hydrolase 1 (UCHL-1) in human myeloma cells. Br J Haematol **2004**;127:292-8

2.	Ogawa A, Izumikawa K, Tate S, Isoyama S, Mori M, Fujiwara K*, et al.* SLFN11-mediated ribosome biogenesis impairment induces TP53-independent apoptosis. Mol Cell **2025**;85:894-912 e10

3.	Kim JM, Kee Y, Gurtan A, D'Andrea AD. Cell cycle-dependent chromatin loading of the Fanconi anemia core complex by FANCM/FAAP24. Blood **2008**;111:5215-22

4.	Murai J, Feng Y, Yu GK, Ru Y, Tang SW, Shen Y*, et al.* Resistance to PARP inhibitors by SLFN11 inactivation can be overcome by ATR inhibition. Oncotarget **2016**;7:76534-50

5.	Ikeda S, Kitadate A, Abe F, Takahashi N, Tagawa H. Hypoxia-inducible KDM3A addiction in multiple myeloma. Blood Adv **2018**;2:323-34

6.	Caicedo HH, Hashimoto DA, Caicedo JC, Pentland A, Pisano GP. Overcoming barriers to early disease intervention. Nat Biotechnol **2020**;38:669-73

7.	Goldman MJ, Craft B, Hastie M, Repecka K, McDade F, Kamath A*, et al.* Visualizing and interpreting cancer genomics data via the Xena platform. Nat Biotechnol **2020**;38:675-8

8.	Zhan F, Huang Y, Colla S, Stewart JP, Hanamura I, Gupta S*, et al.* The molecular classification of multiple myeloma. Blood **2006**;108:2020-8

9.	Zhan F, Barlogie B, Arzoumanian V, Huang Y, Williams DR, Hollmig K*, et al.* Gene-expression signature of benign monoclonal gammopathy evident in multiple myeloma is linked to good prognosis. Blood **2007**;109:1692-700

10.	Luna A, Elloumi F, Varma S, Wang Y, Rajapakse VN, Aladjem MI*, et al.* CellMiner Cross-Database (CellMinerCDB) version 1.2: Exploration of patient-derived cancer cell line pharmacogenomics. Nucleic Acids Res **2021**;49:D1083-D93

11.	Broyl A, Hose D, Lokhorst H, de Knegt Y, Peeters J, Jauch A*, et al.* Gene expression profiling for molecular classification of multiple myeloma in newly diagnosed patients. Blood **2010**;116:2543-53

12.	Ubels J, Sonneveld P, van Beers EH, Broijl A, van Vliet MH, de Ridder J. Predicting treatment benefit in multiple myeloma through simulation of alternative treatment effects. Nat Commun **2018**;9:2943

13.	Law CW, Chen Y, Shi W, Smyth GK. voom: Precision weights unlock linear model analysis tools for RNA-seq read counts. Genome Biol **2014**;15:R29

14.	Ritchie ME, Phipson B, Wu D, Hu Y, Law CW, Shi W*, et al.* limma powers differential expression analyses for RNA-sequencing and microarray studies. Nucleic Acids Res **2015**;43:e47

15.	Ashburner M, Ball CA, Blake JA, Botstein D, Butler H, Cherry JM*, et al.* Gene ontology: tool for the unification of biology. The Gene Ontology Consortium. Nat Genet **2000**;25:25-9

16.	Kuleshov MV, Jones MR, Rouillard AD, Fernandez NF, Duan Q, Wang Z*, et al.* Enrichr: a comprehensive gene set enrichment analysis web server 2016 update. Nucleic Acids Res **2016**;44:W90-7

17.	Lachmann A, Xie Z, Ma'ayan A. blitzGSEA: efficient computation of gene set enrichment analysis through gamma distribution approximation. Bioinformatics **2022**;38:2356-7

**Supplemental Figure Legends**

**Figure S1. *SLFN11* expression in cancer cell lines and multiple myeloma subtypes. (A)** Scatter plot showing the relationship between *SLFN11* and *MKI67* (RNA-seq expression) across the 1,019 cancer cell lines in the CCLE-Broad-MIT database. Multiple myeloma cell lines (N=27, red) and other cancer cell lines (grey) are shown (data accessed through CellMinerCDB (<https://discover.nci.nih.gov/cellminercdb/>). (B) *SLFN11* expression levels (log2(fpkm-uq+1)) across subtypes of multiple myeloma as defined by Maura et al. (Nature Genetics, 2024). Subtypes include PR, 1q gain, MS, HRD with various genomic alterations (++15, MYC, low TP53, low NFκB), CD1, CD2a, CD2b, MAF/MAFB, and low purity samples. Bars represent mean values. **(C)** SLFN11 expression levels across copy number-based subtypes of multiple myeloma as defined in the same study. Subtypes include +1q/-13, -13, diploid, and HRD with various ploidy patterns. Bars represent mean values.

**Figure S2. SLFN11/CD138 expression patterns and PCPI activity in sequential MM samples. (A)** High-magnification image of dual immunohistochemical staining for CD138 (blue, membrane) and SLFN11 (brown, nuclear) showing four distinct cell populations. Purple arrowheads indicate CD138+/SLFN11+ cells, blue arrowheads CD138+/SLFN11- cells, brown arrowheads CD138-/SLFN11+ cells, and white arrowheads CD138-/SLFN11- cells. Scale bar: 10 μm. (B) Quantitative analysis of the 4 cell populations in sequential bone marrow samples from three MM patients (Patients 1-3 from Figure 2) at primary diagnosis and through multiple relapses. Normal bone marrow controls are shown for comparison. Cell populations were quantified by counting 1,000 cells per sample (or 500 cells for low-density samples). (C) Plasma Cell Proliferation Index (PCPI) showing progressive increases with each relapse in the 3 patients examined. PCPI was calculated as the percentage of CD138-positive cells co-expressing Ki-67 by counting 200 CD138-positive cells per sample. Normal bone marrow samples show minimal PCPI values.

**Figure S3. Correlation of *SLFN11* expression with lineage markers and transcription factors in multiple myeloma. (A)** Scatter plots showing lack of correlation between SLFN11 expression and non-plasma cell lineage markers in multiple myeloma samples (n = 844): *CD8A* (T-cell marker, r = -0.038), *CD79A* (B-cell marker, r = -0.0052), and *CD68* (monocyte/macrophage marker, r = -0.090), contrasting with the positive correlations observed with plasma cell markers in Figure 3C-E. (B) Heatmap showing correlation between *SLFN11* expression and all 122 transcription factors and epigenetic enzymes involved in plasma cell differentiation across 844 multiple myeloma samples, expanding on the top correlations shown in Figure 3D. (C) Scatter plot showing the correlation between *SLFN11* and *FLI1* (r = 0.15, P < 1 × 10^-15^), a known transcriptional regulator of *SLFN11* expression.

**Figure S4. Heatmap of pathway-related genes and comprehensive GSEA analysis. (A)** Heatmap showing expression patterns of differentially expressed genes across 844 multiple myeloma samples ordered by *SLFN11* expression (high to low). Genes are grouped by functional pathways: ER stress response, ERAD (ER-associated degradation), ubiquitin-protein ligase, ubiquitin-transferase, and ribosome. Red indicates high expression; blue indicates low expression. Gene lists for each pathway are provided in Supplemental Table S1. **(B)** Comprehensive Gene Set Enrichment Analysis (GSEA) results comparing *SLFN11*-high versus *SLFN11*-low multiple myeloma samples (n = 844, divided by median expression). Left panel: MSigDB Hallmark gene sets. Right panel: MSigDB C5 ontology gene sets. Red indicates positive enrichment in *SLFN11*-high samples; blue indicates negative enrichment (positive in *SLFN11*-low samples). Selected representative enrichment plots are shown in Figure 4B. NES: Normalized Enrichment Score; SIZE: gene set size.

**Figure S5. SLFN11 recruitment to nucleoli occurs in doxycycline-inducible SLFN11-expressing U2OS cells.** (A) Representative immunofluorescence images showing chromatin-bound SLFN11 localization (magenta, upper panels) and merged images with DAPI (blue, lower panels) in doxycycline-induced U2OS cells. Cells were treated with camptothecin (CPT, positive control for chromatin-bound SLFN11), or bortezomib (BTZ, 1 µM) for 4 hours followed by pre-extraction, fixation, and staining. CPT treatment shows SLFN11 localization at replication forks, while BTZ treatment induces SLFN11 translocation to nucleoli. Scale bar: 10 µm. (B) Quantification of SLFN11 signal intensities in individual cells (n = 2453-2927 cells per condition). Mean ± SD are shown. ***P < 0.001, ****P < 0.0001 (one-way ANOVA). a.u., arbitrary units. (C) Quantifications of the proportion of cells with chromatin-bound SLFN11 foci overlapping with nucleolin in Figure 5D. Mean ± SD are shown. *P < 0.05, **P < 0.01 (one-way ANOVA).

**Figure S6. SLFN11 recruitment to nucleoli limits ribosomal RNA (rRNA) synthesis and affects global translation rates after Bortezomib treatment. (A)** Treatment protocol. Doxycycline (DOX)-inducible SLFN11-expressing U2OS cells were treated with bortezomib (BTZ, 1 μM) for 4 hours. **(B)** Representative immunofluorescence images showing 5-ethynyluridine (EU) incorporation (green). Scale bar: 10 µm. (C) Quantification of EU signals in individual cells for the indicated treatments (n = 135 – 167 cells per condition). Mean ± SEM are shown. ****P < 0.0001 (one-way ANOVA). a.u., arbitrary units. The threshold for EU (70) was determined based on distributions observed in the control experiment. a.u., arbitrary units. (D) Representative immunofluorescence images showing HPG incorporation (green). WT and SLFN11 KO clones of KMS-27 cells were treated with BTZ (0.1 µM) for indicated timepoints. Scale bar: 10 µm. (E) Quantification of HPG signals in individual cells for the indicated treatments (n = 120 – 159 cells per condition). Mean ± SEM are shown. ****P < 0.0001 (one-way ANOVA). a.u., arbitrary units.

**Figure S7. Event-free survival by treatment arm stratified by SLFN11 expression in the HOVON-65/GMMG-HD4 trial.** Kaplan-Meier analyses comparing PAD versus VAD within each SLFN11 expression subgroup (unadjusted log-rank test). **(A)** SLFN11-low patients (n=163; PAD, n=84; VAD, n=79; log-rank P=0.072). **(B)** SLFN11-high patients (n=164; PAD, n=85; VAD, n=79; log-rank P=0.89). ISS stage-adjusted analyses incorporating both subgroups are presented in Figure 8C. EFS, event-free survival; PAD, bortezomib/doxorubicin/dexamethasone; VAD, vincristine/doxorubicin/dexamethasone.
