## Supplementary figures and images for "Schlafen 11 (SLFN11) overexpression in multiple myeloma and nucleolar translocation in response to bortezomib"

### Figure S1. SLFN11 expression

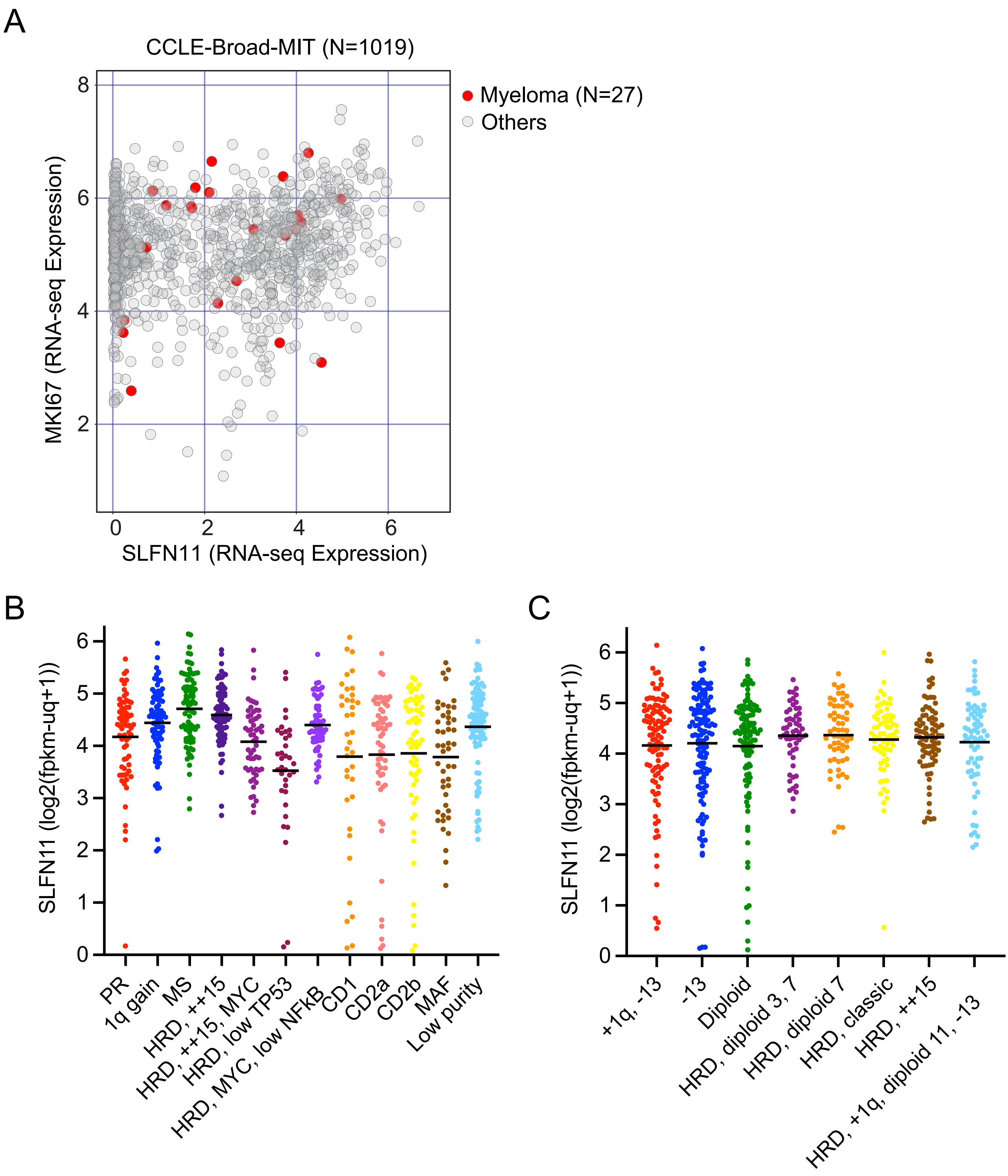

### Figure S2. SLFN11/CD138 patterns

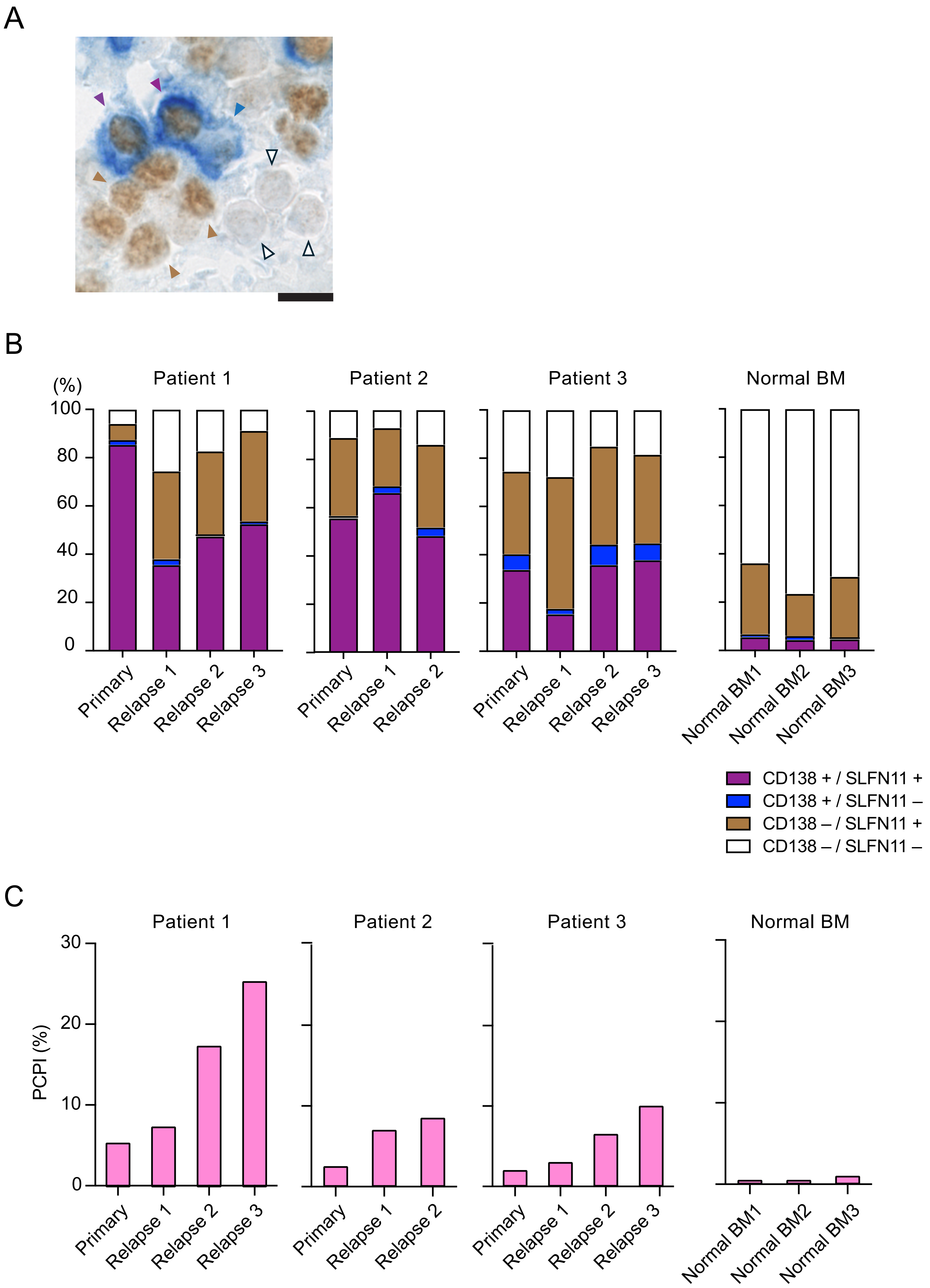

### Figure S3. SLFN11 correlations

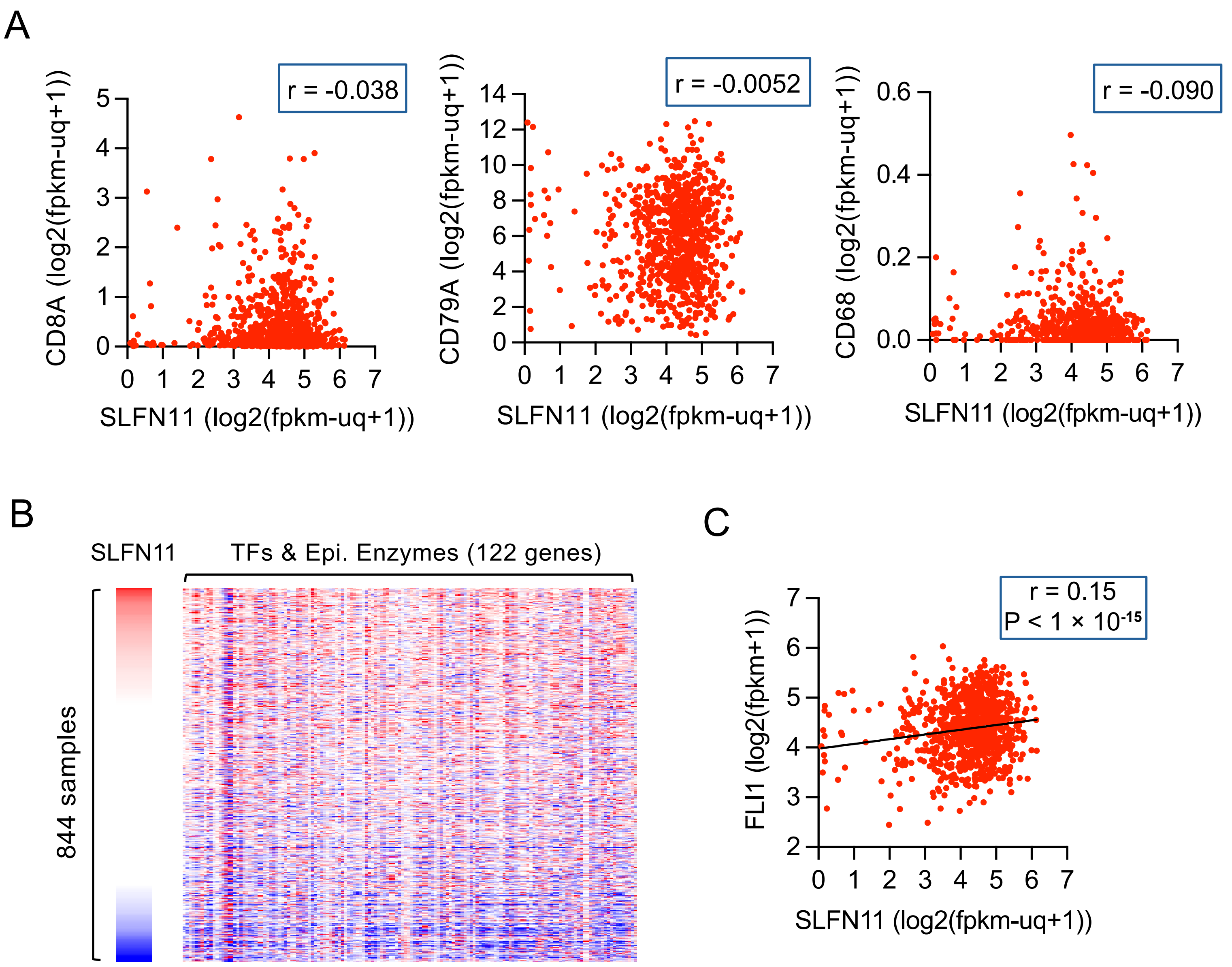

### Figure S5. SLFN11 in U2OS

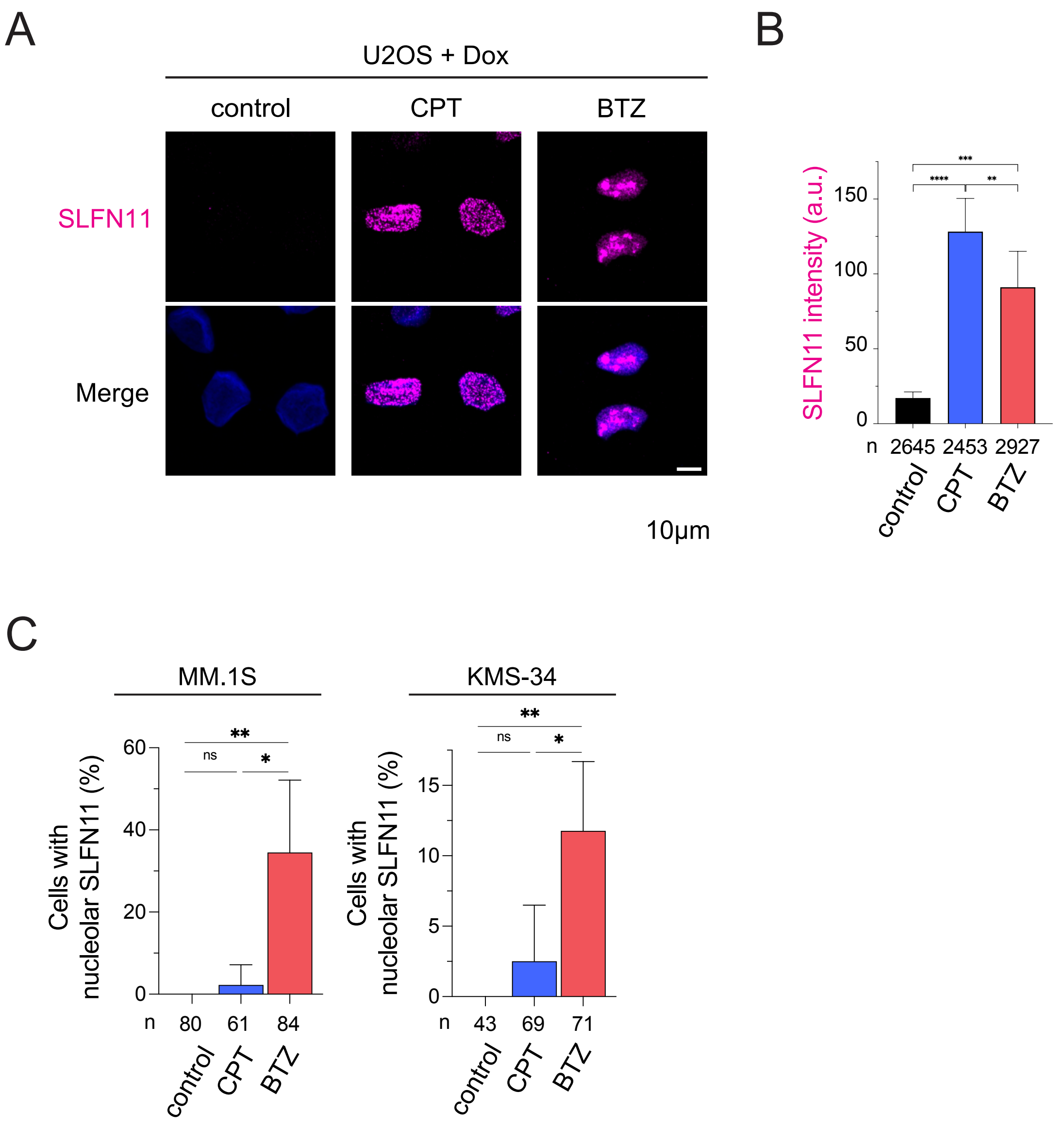

### Figure S6. rRNA and translation

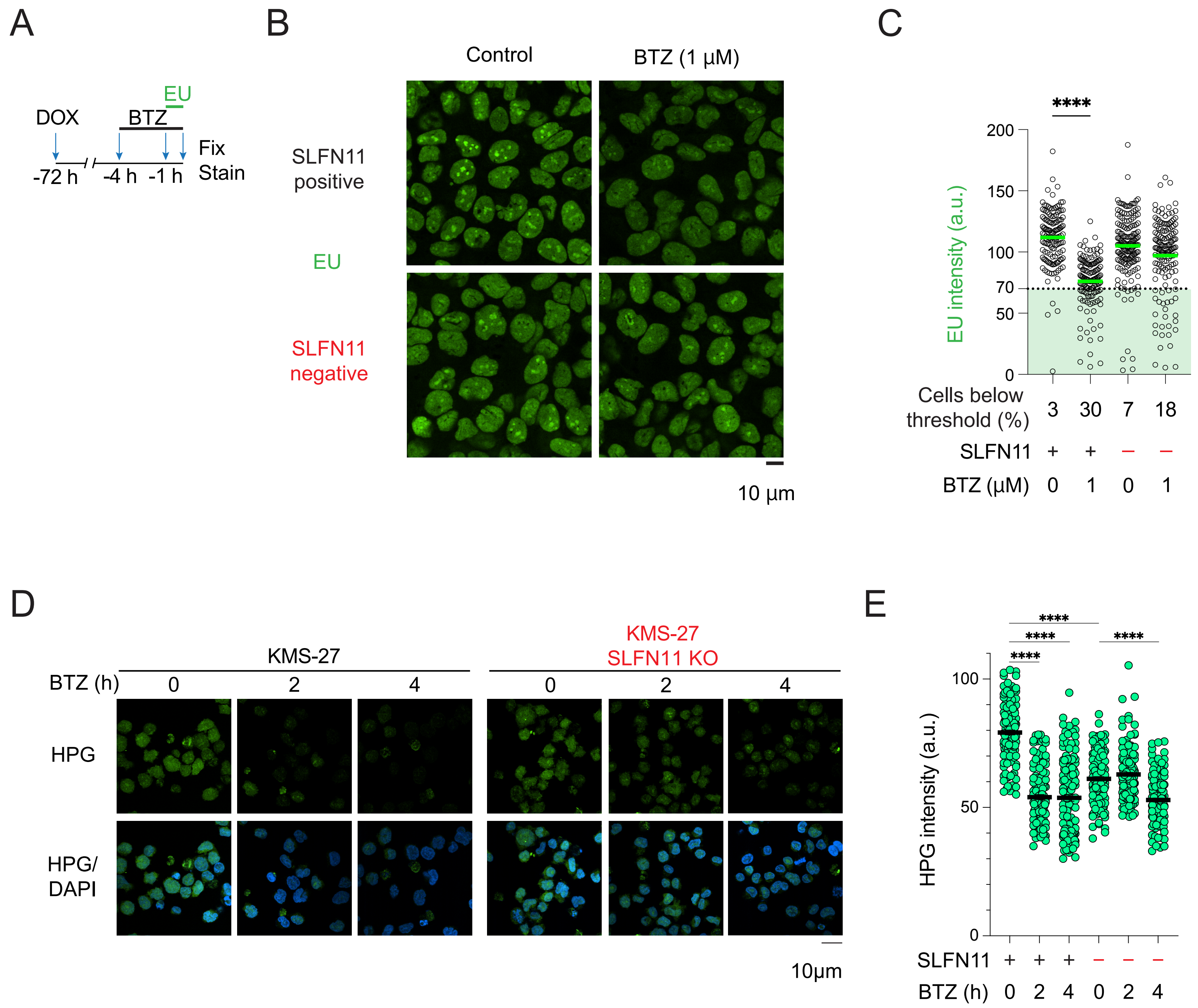

### Figure S7. HOVON-65 EFS

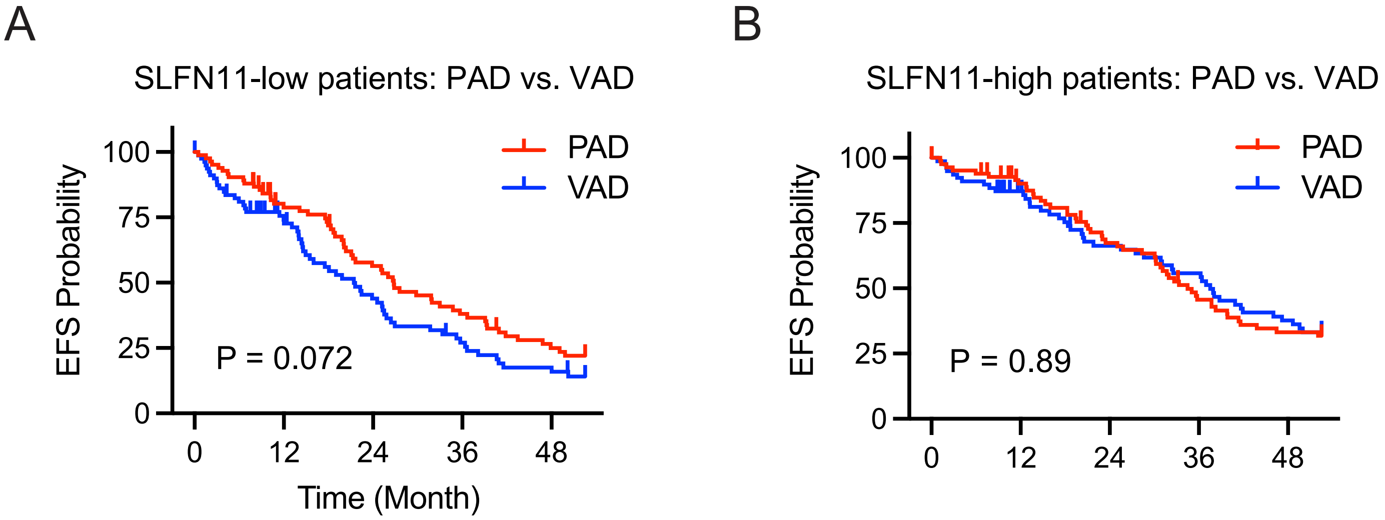
