## Supplementary material for "Schlafen 11 (SLFN11) overexpression in multiple myeloma and nucleolar translocation in response to bortezomib": Figure S4. Pathways and GSEA

A

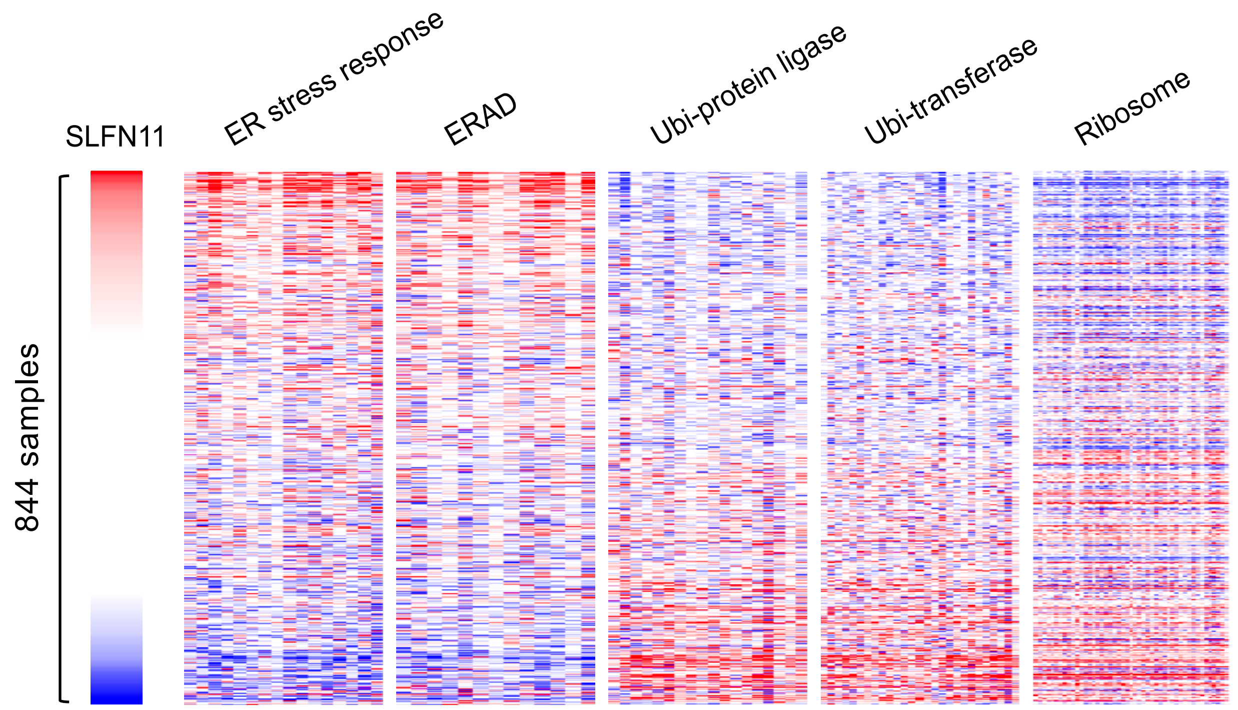


B

| NES | SIZE | SET |
|---|---|---|
| -5.275 | 85 | TNFα signaling via NFκB |
| 3.460 | 84 | Unfolded protein response |
| -3.154 | 48 | Inflammatory response |
| -3.038 | 82 | Apoptosis |
| -2.955 | 92 | P53 pathway |
| -2.757 | 168 | Myc targets v1 |
| 2.643 | 80 | Glycolysis |
| -2.385 | 83 | G2M checkpoint |
| -2.158 | 56 | IL2 STAT5 signaling |
| 2.092 | 62 | Protein secretion |

MSigDB H: hallmark gene sets

| NES | SIZE | SET |
|---|---|---|
| 8.270 | 219 | Adaptive immune response |
| 8.004 | 95 | Immunoglobulin complex |
| 7.679 | 98 | Antigen binding |
| -7.328 | 116 | Cytoplasmic translation |
| 7.112 | 120 | External side of plasma membrane |
| -7.023 | 87 | Cytosolic ribosome |
| 7.017 | 128 | Lymphocyte mediated immunity |
| -6.955 | 130 | Structual constituent of ribosome |
| 6.910 | 202 | Cell surface |
| -6.756 | 158 | Ribosome |

MSigDB C5: ontology gene sets
